# Border-forming wound repair astrocytes

**DOI:** 10.1101/2023.08.25.554857

**Authors:** Timothy M. O’Shea, Yan Ao, Shinong Wang, Yilong Ren, Amy Cheng, Riki Kawaguchi, Vivek Swarup, Michael V. Sofroniew

## Abstract

Central nervous system (CNS) lesions become surrounded by neuroprotective borders of newly proliferated reactive astrocytes. Fundamental features of these cells are poorly understood. Here, we show that 90% of border-forming astrocytes derive from proliferating Aldh1l1-expressing local astrocytes, and 10% from Pdgfra-expressing oligodendrocyte progenitors in mice. Temporal transcriptome analysis, snRNAseq and immunohistochemistry showed that after CNS injury, local mature astrocytes dedifferentiated, proliferated, and became transcriptionally reprogrammed to permanently altered new functional states, with persisting downregulation of molecules associated with astrocyte-neuron interactions, and upregulation of molecules associated with wound healing, microbial defense, and interactions with stromal and immune cells. Our findings show that at CNS injury sites, local mature astrocytes proliferate and adopt canonical features of essential wound repair cells that persist in adaptive states and are the predominant source of neuroprotective borders that re-establish CNS integrity by separating neural parenchyma from stromal and immune cells as occurs throughout the healthy CNS.

## Introduction

All organs share the ability to rapidly repair tissue lesions by generating newly proliferated cells that derive from stromal-, immune-, and parenchymal-cell lineages. This multicellular proliferative wound response is a protective adaptation that limits tissue damage, sustains organ integrity and function, and is essential for organism survival^1^. In the central nervous system (CNS), astrocytes are key components of a multicellular proliferative wound response that is stimulated by tissue damage across a broad cross-section of CNS disorders including traumatic injury, stroke, infection, autoimmune inflammation, and certain neurodegenerative diseases^2–6^.

Understanding the derivation and temporally regulated proliferation, maturation, and functions of newly proliferated astrocytes during this wound response is fundamental to understanding and ameliorating the pathophysiology of many different CNS disorders.

Astrocytes are CNS parenchymal cells of neural progenitor cell origin^7^. They contiguously tile the entire CNS and provide multiple activities essential for CNS function in health and disease^2,8–12^. Astrocytes rarely divide in healthy adult CNS^13^ and exist in a state of potentially reversible cell cycle arrest (G0) ^14^. Astrocytes respond to all forms of CNS injury and disease with molecular, structural, and functional changes commonly referred to as astrocyte reactivity^2,6^. Notably, astrocyte reactivity can either be non-proliferative or proliferative^2,13,15^ and is tailored to different disorder contexts by multifactorial signaling mechanisms^16^.

Proliferative astrocyte reactivity occurs in response to overt CNS tissue damage and results in the formation of borders that surround tissue damaged by trauma, ischemia, infection, autoimmune inflammation, fibrosis, neoplasm, foreign bodies or pronounced neurodegeneration^4,13,15,17–24^. Multiple genetically targeted loss-of-function studies demonstrate that newly proliferated astrocyte borders serve essential functions that protect adjacent viable neural tissue, such that transgenic ablation or attenuation of border-forming astrocytes leads to impaired neural parenchymal wound repair with greater spread of destructive inflammation, larger fibrotic lesions, increased loss of neural tissue and impairment of neurological recovery ^4,13,16,17,22,23,25–28^. Despite the increasingly recognized importance of newly proliferated border-forming astrocytes, fundamental features of these cells are poorly understood. Here, we used a combination of transgenically targeted lineage tracing^29,30^, astrocyte specific transcriptome analysis^16,31^, single nucleus RNA sequencing and immunohistochemical protein detection in mice to identify and selectively profile the transcriptional changes over time after injury of the predominant cellular source of newly proliferated astrocytes after traumatic injury.

## Results

### Derivation of lesion border astrocytes

Studies from multiple laboratories implicate two main potential cellular sources for newly proliferated astrocytes around CNS injuries: local astrocytes^17–19,25,32^, and local oligodendrocyte progenitor cells (OPC) ^32–35^. Here, we used lineage-tracing based on tamoxifen regulated Cre-reporter expression^29^, to determine the proportional contributions of these cell types to newly proliferated border-forming astrocytes around hemorrhagic lesions after crush SCI or around ischemic lesions after forebrain stroke caused by infusion of L-NIO (Fig. 1, Extended Data Fig. 1). We targeted the reporter, tdT, to mature astrocytes by using Aldh1l1-CreERT^36^, and to OPC by using either Pdgfra-CreERT-tdT^37^ or NG2-CreERT-tdT^38,39^ and induced temporary Cre expression with a 5 day regimen of tamoxifen dosing in healthy young adult (>8 week old) mice (Fig. 1a).

**Fig. 1.**
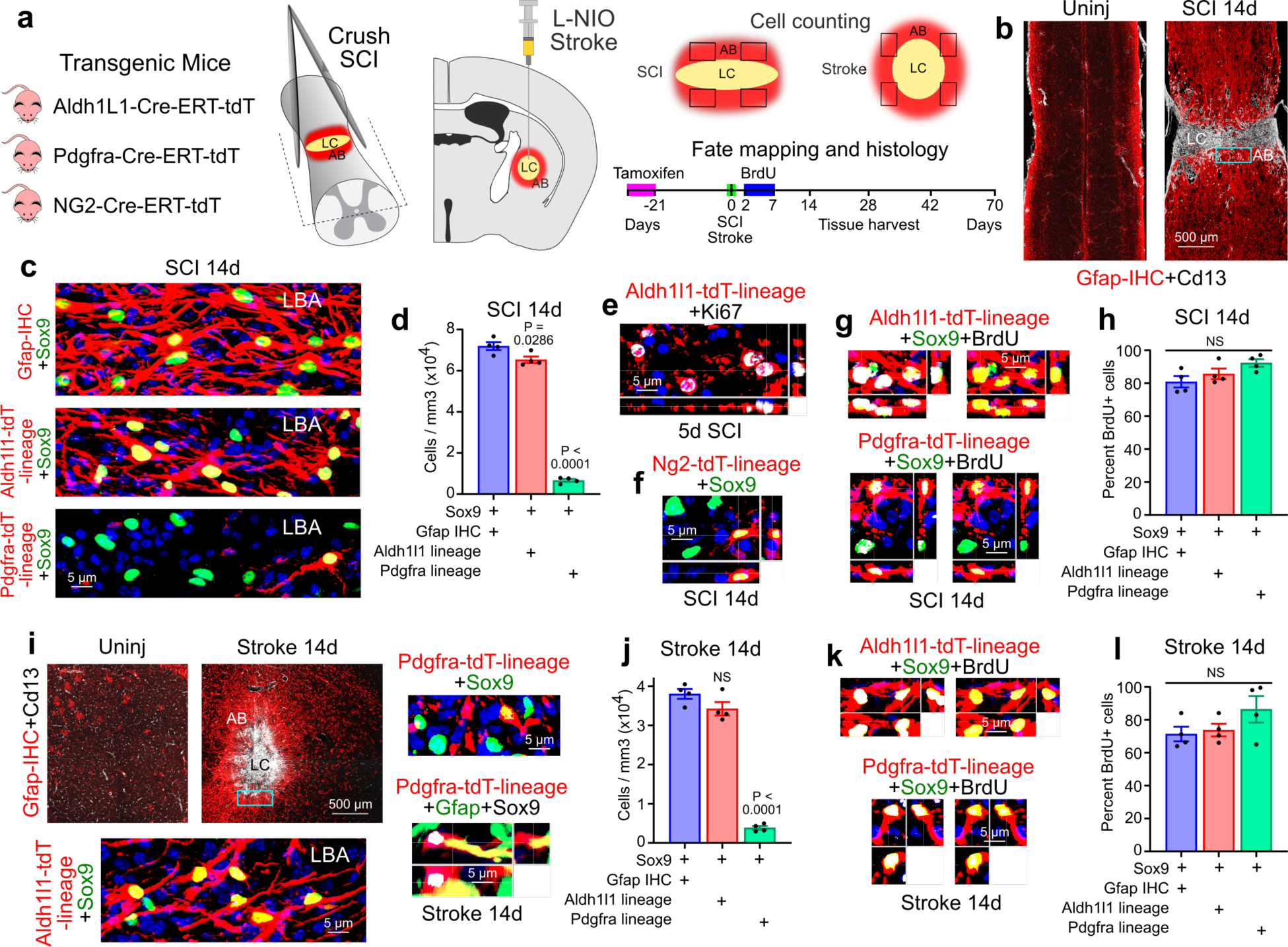
Lineage of border-forming astrocytes that surround CNS lesions. **a.** Lineage tracing procedures. **b.** Spinal cord uninjured and after SCI stained by immunohistochemistry (IHC) for astrocytes (Gfap) or stromal cells (Cd13). **c,d.** Images (c) and cell counts (d) of Sox9-positive lesion border astrocytes (LBA) plus Gfap-IHC or of lineage tracing with Aldh1l1-tdT or Pdgfra-tdT after SCI. **e.** Proliferating astrocytes labeled with Ki67. **f.** Staining for Sox9 plus Ng2-tdT. **g,h.** Newly-proliferated BrdU-labelled astrocytes positive for Aldh1l1-tdT or Pdgfra-tdT after SCI (individual fluorescence channels are shown in Extended Data Figure 1g). **i,j.** Striatum uninjured and after L-NIO stroke, with images (i) and cell counts (j) of Sox9-positive LBA plus Gfap-IHC or of lineage tracing with Aldh1l1-tdT or Pdgfra-tdT after stroke. **k,l.** Newly-proliferated BrdU-labelled astrocytes positive for Aldh1l1-tdT or Pdgfra-tdT after stroke. n = 4 for all groups; ANOVA with Tuckey’s post hoc comparison. LC lesion core, AB astrocyte border

Newly proliferated astrocyte borders around CNS lesions mature into narrow zones of astrocytes in high cell density with overlapping cell processes that surround non-neural lesion cores of stromal and fibrotic tissue after SCI or stroke and are essentially permanent (Fig. 1b,c,i; Extended Data Fig. 1a). We quantified tdT-labelled, lineage traced, border-forming astrocytes within 250µm zones immediately adjacent to lesion core stromal tissue at 14 days after SCI or forebrain stroke (Fig. 1a-c,i,j), a time point by which border formation is largely complete^13^. As benchmarks against which to quantitatively compare tdT labelling, we used Gfap and Sox9, which together label essentially all newly-proliferated border-forming astrocytes around lesions^13,25,30^ (Fig. 1c,d,i,j).

In healthy adult spinal cord or striatum, essentially all astrocytes expressed Sox9 and Aldh1l1-CreERT-tdT, confirming previous reports^36^ and no astrocytes were detectably derived from OPC as indicated by Pdgfra lineage tracing (Extended Data Fig. 1b). In uninjured spinal cord, all Sox9 and Aldh1l1-tdT positive astrocytes expressed detectable Gfap, whereas in uninjured striatum only about 13% did so (Extended Data Fig. 1b).

Lineage tracing showed that after both SCI and stroke, over 90% of Gfap plus Sox9-positive lesion border astrocytes also expressed Aldh1l1-CreERT-tdT, indicating that these cells derived from local mature astrocytes (Fig. 1c,d,i,j). Approximately 10% of Gfap plus Sox9-positive lesion border astrocytes were derived from Pdgfra-CreERT-tdT progeny and also expressed transcription factors Sox10 and Id3, indicating that these cells descended from local OPC and exhibited molecular features of reactive astrocytes^16,40^ (Fig. 1c,d,f,i,j; Extended Data Fig. 1c-e). The proportion of OPC derived border forming astrocytes was essentially equivalent using NG2-CreERT-tdT lineage tracing (Fig. 1f; Extended Data Fig. 1c,d). In both SCI and stroke, about 75% of the Pdgfra-CreERT-tdT-positive cells in the lesion border zone were Olig2-postive but Gfap- and Sox9-negative, indicating that most of these cells were OPC (Extended Data Fig. 1f).

To identify newly proliferated cells, BrdU was administered during a six-day period from two to seven days after injury (Fig. 1a). In healthy adult spinal cord or striatum, no astrocytes were detectably BrdU-labelled during an equivalent six-day period of BrdU administration whereas about 8 to 10% of OPC were BrdU-labelled during the same period (Extended Data Fig. 1g). At five days after injuries, lineage traced astrocytes expressed the active proliferation marker, Ki67 (Fig. 1e), and quantification of BrdU showed that after both SCI or stroke, at least 75 to 85%, of lesion border astrocytes that were derived from either astrocytes or OPC are newly proliferated (Fig. 1g,h,k,l; Extended Data Fig. 1h) and this is likely a conservative estimate because BrdU was administered only once daily.

### Injury-induced transcriptional reprogramming

Since mature Aldh1l1-expressing astrocytes are the predominant source of border forming cells, we next characterized the temporally dependent transcriptional changes in these astrocytes after SCI. Young adult (3 to 4 months) male and female Aldh1l1-CreERT-RiboTag mice were used to enable Hemagglutinin(HA)-positive ribosome immunoprecipitation (IP) and cell specific transcriptome profiling of astrocytes in heathy uninjured spinal cord and at 2, 5, 14, 28, 42 and 70 days after SCI (Fig. 2a) spanning periods of proliferation, border formation, and chronic border persistence^13,16,24^. Sequencing of mRNA from the flow-through solution after RiboTag IP was used to characterize gene expression by other local cells present in healthy tissue or SCI lesions and enabled assessment of astrocyte enrichment relative to other cells (Extended Data Fig. 2a-c). Aldh1l1-CreERT-RiboTag gave equivalent astrocyte-enriched transcriptional profiles as mGfap-Cre-RiboTag^16,26^ but showed depleted genetic signatures originating from ependyma or OPC-derived cells that acquire Gfap expression after SCI; and both ependyma and OPC had immunohistochemically detectable HA expression in mGfap-Cre-RiboTag but not in Aldh1l1-CreERT-RiboTag spinal cords (Extended Data Fig. 2a-k). Consistent with previous reports after stroke^41^, we detected negligible sex-dependent transcriptomic differences in astrocytes after SCI, with four Y- and two X-chromosome genes detected as the only differentially expressed genes (DEGs) (FDR <0.01) across two independent post SCI time points comparing eight female and eight male mice (Extended Data Fig. 2l-n).

**Fig. 2.**
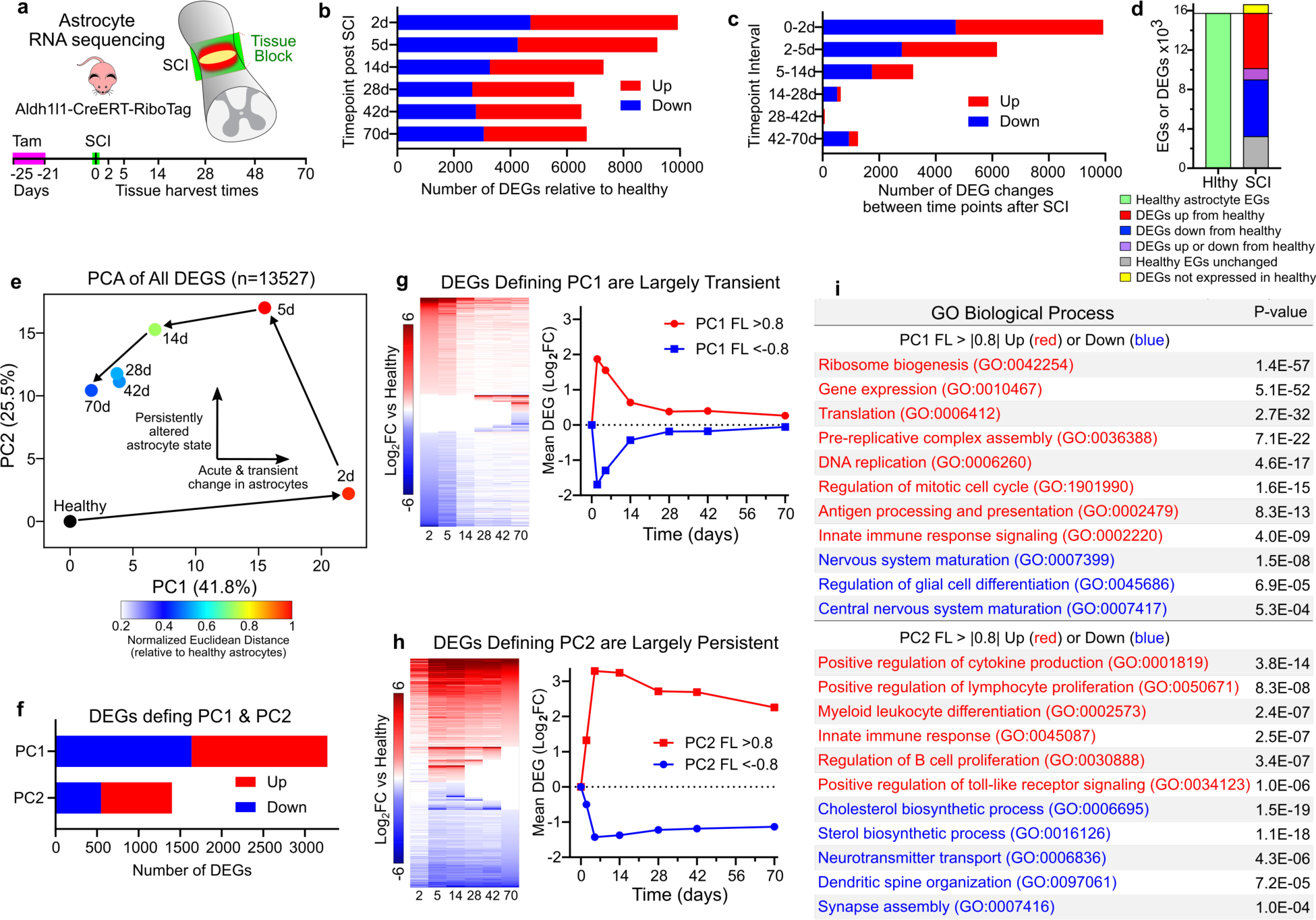
Temporal analysis of SCI-induced astrocyte transcriptional changes. **a.** Experimental design. **b.** Numbers of astrocyte DEGs significantly different (up or down) (FDR<0.01) from uninjured healthy at different times. **c.** Numbers of astrocyte DEG significantly (FDR<0.01) changed between different time points. **d.** Number of healthy astrocyte expressed genes (EGs) compared with DEGs after SCI that are up, down, or not regulated from healthy, or not expressed in healthy. **e.** Principal component analysis (PCA) of all DEGs. **f.** Numbers of DEGs defining PC1 or PC2. **g,h.** Heatmaps and mean DEG log2 fold changes (Log_2_FC) of DEGs defining PC1 (g) and PC2 (h). **i.** GO Biological Processes significantly up (red) or down (blue) regulated as identified by unbiased evaluations of DEGs defined by PC1 or PC2. FL, PC factor loading.

Using Aldh1l1-CreERT-RiboTag (Astro-RiboTag) across all six post SCI timepoints examined, a total of 13,527 unique DEGs were identified compared to baseline levels in healthy astrocytes, with more genes upregulated than downregulated at all timepoints (Fig. 2b). The most DEGs, 9924, were detected at 2 days, with over 6000 DEGs persisting at 28, 42, and 70 days (Fig. 2b). The greatest changes occurred between 0 to 2 and 2 to 5 days, with far fewer changes after 14 days (Fig. 2c).

Over 15,722 genes were identified as expressed by healthy spinal cord astrocytes, and of these, 37% (5755) were primarily downregulated after SCI, 36% (5598) were primarily upregulated, 7% (1153) were dynamically regulated down or up at different timepoints, and 20% (3216) were not significantly different at any timepoint examined (Fig. 2d). Thus, 92% (12,506 of 13,527) of the total DEGs identified across all times after SCI were expressed by astrocytes in healthy tissue (Fig. 2d). Notably, induction of newly expressed genes that were not detectable in healthy astrocytes accounted for only 12% (889 of 7640) of DEGs upregulated by astrocytes after SCI (Fig. 2d).

Principal component analysis (PCA) applied to the entire cohort of 13,527 DEGs identified across all timepoints, revealed a clear temporal progression of astrocyte transcriptional responses with 2 principal components that accounted for over two-thirds of the total system variation (Fig. 2e). PC1, defining the most dominant effect, revealed acute and transient changes. DEGs with a PC1 factor loading of >|0.8| in the positive or negative direction peaked at 2 days after SCI before returning essentially to baseline healthy astrocyte levels by 28 days with roughly equal numbers transiently up- or down-regulated (Fig. 2f,g). DEGs defining PC2 increased quickly and largely persisted with mean values of greater than 2, or less than −1, across the entire time course with more upregulated (854) than downregulated (547) (Fig. 2f,h).

Unsupervised analysis of Gene Ontology (GO) Biological Processes showed that the 1635 transiently upregulated DEGs defining PC1 were enriched for genes associated with gene expression and translational, cell proliferation, innate immune signaling and antigen presentation; whereas the 1636 transiently downregulated DEGs defining PC1 were enriched for genes associated with mature CNS structure and glia cell differentiation (Fig. 2i). The 854 persistently upregulated DEGs defining PC2 were enriched for genes associated with cytokine production, innate and adaptive immune regulation, and extracellular matrix organization, whereas the 547 persistently downregulated DEGs defining PC2 were enriched for genes associated with cholesterol and lipid metabolism, neurotransmitter transport and synapse organization (Fig. 2i).

These findings demonstrate that local healthy mature Aldh1l1-expressing astrocytes undergo pronounced, temporally dependent transcriptional changes during border formation after SCI, including both transient and persisting changes. Remarkably, 88% of DEGs upregulated by astrocytes after SCI were already expressed at detectable levels by healthy astrocytes. Astrocyte border formation involves permanent transcriptional reprogramming since almost 50% of total DEGs persist at 70 days post SCI, including many genes not detectably expressed by healthy astrocytes. By using GO analyses, astrocyte transcriptional changes could be broadly parsed into overlapping profiles related to astrocyte dedifferentiation and proliferation, astrocyte reactivity, regulation of inflammation and immune signaling, wound healing, and persisting border formation, examined in more detail below.

### Astrocyte dedifferentiation, proliferation, and loss-of-functions

We next defined more precisely how local mature astrocytes change in response to SCI. Of the 15,722 total genes expressed by healthy astrocytes (Fig. 2d), about 60% were up- or down-regulated at 2 and 5 days, and about 35% to 40% from 28 through 70 days, with the rest remaining unchanged (Fig. 3a).

**Fig. 3.**
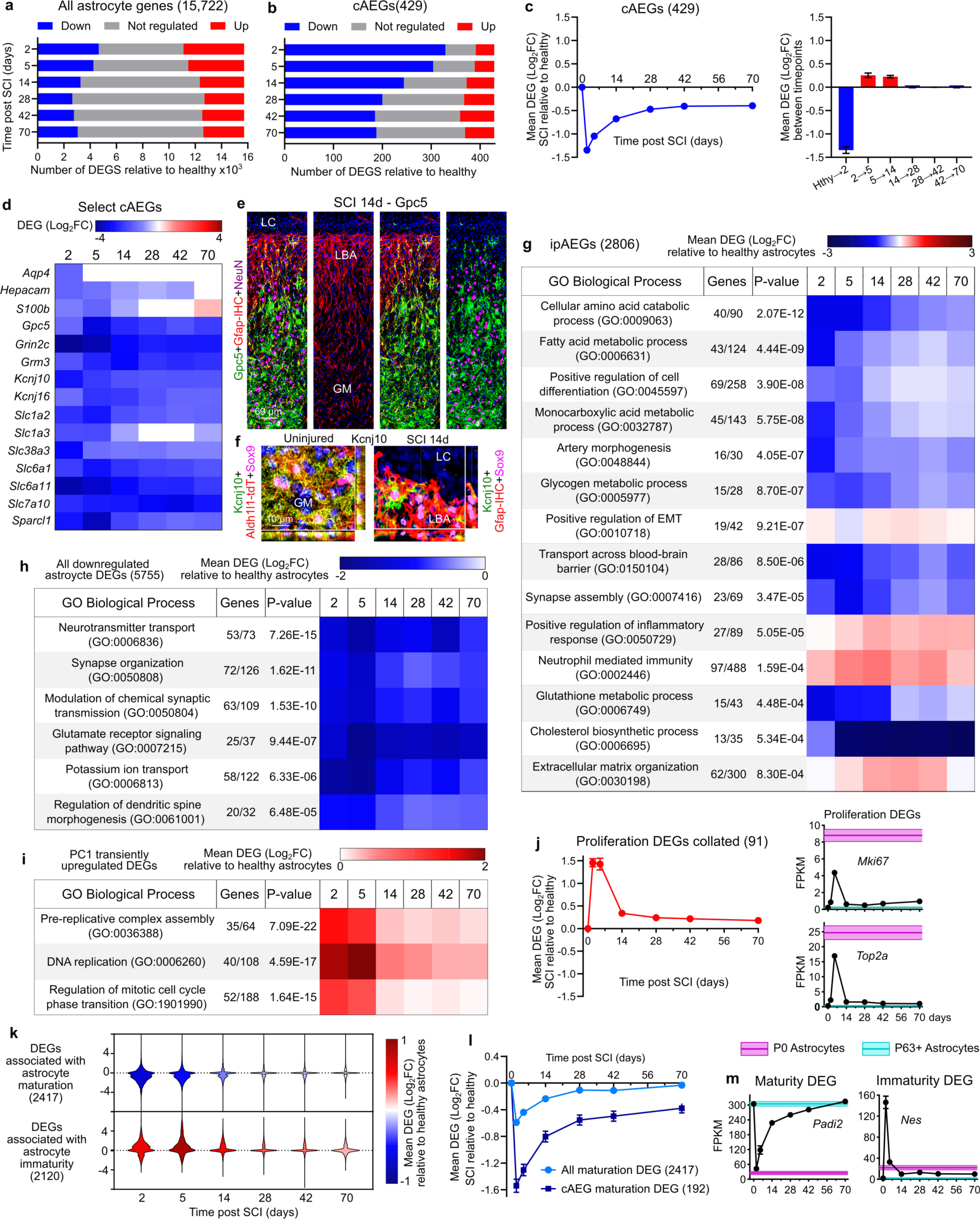
SCI-induced astrocyte dedifferentiation and proliferation. **a.** Numbers of up, down or non-significant changes (FDR <0.01) in 15,722 genes expressed by uninjured astrocytes at different times after SCI. **b.** Numbers of up, down or non-significant changes (FDR <0.01) in 429 consensus astrocyte-enriched genes (cAEG) at different times after SCI. **c.** Mean log2 fold changes (log_2_FC) of down-regulated cAEGs at different times after SCI relative to uninjured; and changes between individual timepoints. **d.** Heatmap of mean log_2_FC of selected down-regulated cAEGs at different times after SCI. **e,f.** Markedly reduced Gpc5 (e) and Kcnj10 (f) immunoreactivities in lesion border astrocytes (LBA) compared with more distal grey matter (GM) astrocytes. For individual fluorescence channels see Extended Data Fig. 3c. **g.** Time course of mean expression changes after SCI of ipAEGs (see main text) associated with representative examples of the most significantly changed GO Biological Processes associated with ipAEGs. **h.** Highly downregulated GO Biological Processes associated with all downregulated astrocyte DEGs (whether enriched versus other cells or not). **i.** Time course after SCI of mean changes of DEGs associated with cell proliferation related GO Biological Processes identified by PC1 in Fig. 2i. **j.** Time course after SCI of mean changes in all 91 proliferation-related astrocyte DEGs examined in **i**, and two specific examples, *Mki67* and *Top2a*. **k.** Time courses after SCI of mean changes in consensus genes associated with astrocyte maturity or immaturity. **l.** Time courses after SCI comparing downregulation of all maturation associated genes versus consensus genes expressed by mature astrocytes (cAEGs). **m.** Time courses after SCI of mean changes in two specific examples of DEGs associated with astrocyte maturity or immaturity, *Padi2* and *Nes,* respectively.

To begin characterizing astrocyte-enriched genes, we first examined a panel of 429 consensus healthy astrocyte enriched-genes (cAEGs) identified in at least 5 of 8 published archival datasets^40^, which we confirmed as enriched in our samples of healthy astrocytes by at least 2-fold and up to over 50-fold (Extended Data Fig. 3a,b). Nearly all cAEGs, 97% (417 of 429) were DEGs and only 3% (12 of 429) were not significantly different at any timepoint examined after SCI. Remarkably, 74% (317 of 429) were primarily downregulated across different times and only 11% (49 of 429) were primarily upregulated, with 15% changing in either direction at different times (Fig. 3b). These changes were reflected in a pronounced decrease in the overall mean expression levels of the 429 cAEGs, which had a negative peak at 2 days and was followed by a return towards baseline by 14 and 28 days, but with an overall downregulation persisting through 70 days (Fig. 3c). Notably, cAEGs with the highest enrichment in healthy astrocytes relative to other local cells were the most downregulated (Extended Data Fig. 3b).

Prominent cAEGs that were acutely downregulated and returned to baseline included the water channel, *Aqp4*, the calcium binding protein, *S100b*, and *Hepacam4* which regulates astrocyte branching complexity^42^ (Fig 3d). Prominent cAEGs that were persistently downregulated included transporters for glutamate (*Slc1a2, Slc1a3*), GABA, (*Slc6a1, Slc6a11*), glutamine (*Slc38ac*) and d-serine (*Slc7a10*), potassium channels (*Kcnj10, Kcnj16*), glutamate receptor subunits (*Grin2c, Grm*) and synapse modulating molecules (*Gpc5, Sparcl1*) ^10,43^ (Fig. 3d).

Immunohistochemistry confirmed certain changes at the protein level and showed for example that many, but not all, lesion border astrocytes had low or undetectable levels of Gpc5 and Kcnj10 (Fig. 3e,f; Extended Data Fig. 3c).

To explore more broadly how local healthy astrocytes acutely changed functional states after SCI, we examined several additional DEG cohorts. We first expanded our evaluation of astrocyte enriched genes by examining 2806 genes identified by our RiboTag IP as significantly enriched in healthy astrocytes by at least 2-fold and up to 50-fold greater expression versus other local cells (Extended Data Fig. 3a,b). Similar to cAEGs, these IP astrocyte enriched genes (ipAEGs) exhibited a preponderance of downregulation after SCI, particularly among those ipAEGs most highly enriched in healthy astrocytes (Extended Data Fig. 3b). We conducted unsupervised analysis of GO Biological Processes (GO-BP) associated with significant changes among these 2806 ipAEGs after SCI, and then tracked over time after SCI the mean expression of ipAEGs associated with representative examples of the most significantly changed GO-BP. The majority of significantly altered GO-BP were associated with downregulated ipAEGs related to cell differentiation, fatty acid metabolism, general metabolic processes, vascular morphogenesis, transport across the blood-brain barrier, synapse assembly, glutathione production, and cholesterol production (Fig. 3g). The few GO-BP associated with upregulated ipAEGs were related to epithelial-to-mesenchymal transition (EMT), immune functions and extracellular matrix reorganization.

We next examined GO-BP associated with all 5755 DEGs downregulated by astrocytes at any time after SCI, and again tracked over time after SCI the mean expression of DEGs associated with representative examples of the most significantly changed. This analysis further confirmed a pronounced acute and persistent attenuation of astrocyte functions associated with astrocyte-neuron interactions, neurotransmitter transport, synapse organization, synaptic transmission, and potassium regulation (Fig. 3h).

Cell dedifferentiation can be associated with proliferation^44^. Past^13^ and present Ki67 and BrdU evaluations (Fig. 1e,g,h,k,l) and GO-BP analysis (Fig 2i) indicate pronounced astrocyte proliferation starting around 2 days after SCI. We tracked over time after SCI changes among genes associated with three of the most significantly upregulated cell proliferation related GO-BP identified by PCA (Fig. 2i, 3i,j). Astrocyte DEGs associated with each of these GO-BP, and the mean expression of the 91 proliferation-related genes associated with all three GO-BP, were highly upregulated by local astrocytes at 2 and 5 days and returned to near baseline levels by 14 days (Fig. 3i,j).

Dedifferentiation and proliferation of local healthy Aldh1l1-expressing astrocytes after SCI suggested a potential return to an immature or progenitor-like state^45^. We compared astrocyte DEGs after SCI with two panels of 2120 and 2417 DEGs that were positively or negatively associated with the progression of astrocyte maturation during postnatal development. These panels were derived by PCA of transcriptomes from healthy astrocytes at postnatal days (P) from P0 to P63 (Extended Data Fig. 3d-h). By 2 and 5 days after SCI, astrocytes had markedly downregulated mean expression of 2417 genes associated with maturity and upregulated mean expression of 2120 genes associated with immaturity and these mean changes returned to essentially baseline levels by 28 days, with a modest upregulation of some immaturity genes persisting to 70 days (Fig. 3k; Extended Data Fig. 3d-h). Of the 429 cAEGs, 192 were positively associated with the progression towards maturity and their mean expression declined and remained persistently low after SCI (Fig. 3l,m). Notably, healthy P0 astrocytes expressed high levels of *Mki67* and *Top2a* associated with active proliferation (Fig. 3j), whereas mature astrocytes in healthy CNS are proliferation-dormant and exist in a potentially reversible G0 state (Extended Data Figs. 1g) and required two or more days after injury to express these genes and never reach FPKM levels of healthy proliferating P0 astrocytes (Fig. 3j).

These findings demonstrate that local healthy mature Aldh1l1-expressing astrocytes respond acutely to SCI with downregulation of most astrocyte enriched genes and a transient phase of proliferation and immaturity. This is followed by a return of many transcriptional features of mature astrocytes but with persisting differences, including a persistent down-regulation of molecules associated with astrocyte-neuron interactions such as maintenance of extracellular neurotransmitter and ion homeostasis, and synapse organization and function. These findings point towards persistent transcriptional reprogramming of newly proliferated lesion border astrocytes to new and different functional states.

### Astrocyte reactivity and gain-of-functions

To look for potential new functions adopted by newly proliferated and reprogrammed astrocytes after SCI, we first examined changes in 170 consensus astrocyte reactivity genes (cARGs) derived from a meta-analysis of six archival datasets from multiple laboratories^40^. All 170 cARGs were upregulated on at least one timepoint after SCI and 93% (158 of 170) were upregulated at all timepoints. 92% (157 of 170) were detectably expressed by healthy astrocytes. Mean cARG expression increased 16-fold by 2 days after SCI, increased to over 50-fold by 5 days and then declined moderately but remained persistently elevated by over 8-fold at 70 days (Fig. 4b), consistent with reprogramming to an essentially permanent reactive state after SCI. Notably, 41 of the top 50 GO-BP most significantly associated with cARGs upregulated at all timepoints involved regulation of inflammation, while other upregulated GO-BP included phagocytosis, extracellular matrix organization, synapse pruning and homotypic cell-cell adhesion (Extended Data Fig. 4a). cARGs that were upregulated by 14 days after SCI and remained persistently and highly upregulated at 70 days included well studied cARGs such as *Gfap*, *Vim* and *Lgals3,* as well as molecules that appear in multiple reactive astrocyte RNA evaluation studies such as *S100a6, Serpina3n, Lyz2, Lcn2, Hsbp1, C1qa, Tyrobp, Trem2, Tgm1*, *Ccl3* and *Ccl4* (Fig. 4c). Immunohistochemistry confirmed protein expression by border forming astrocytes for many of these molecules, including after stroke (Fig. 4d, Extended Data Fig. 4b). Notably, immunohistochemistry revealed that whereas certain proteins such as Gfap and S100a6 were readily detectable in essentially all Sox9-positive lesion border astrocytes, many proteins such as Lgals3 and others discussed below, were detectably expressed in some but not all lesion border astrocytes (Extended Data Fig. 4b).

**Fig. 4.**
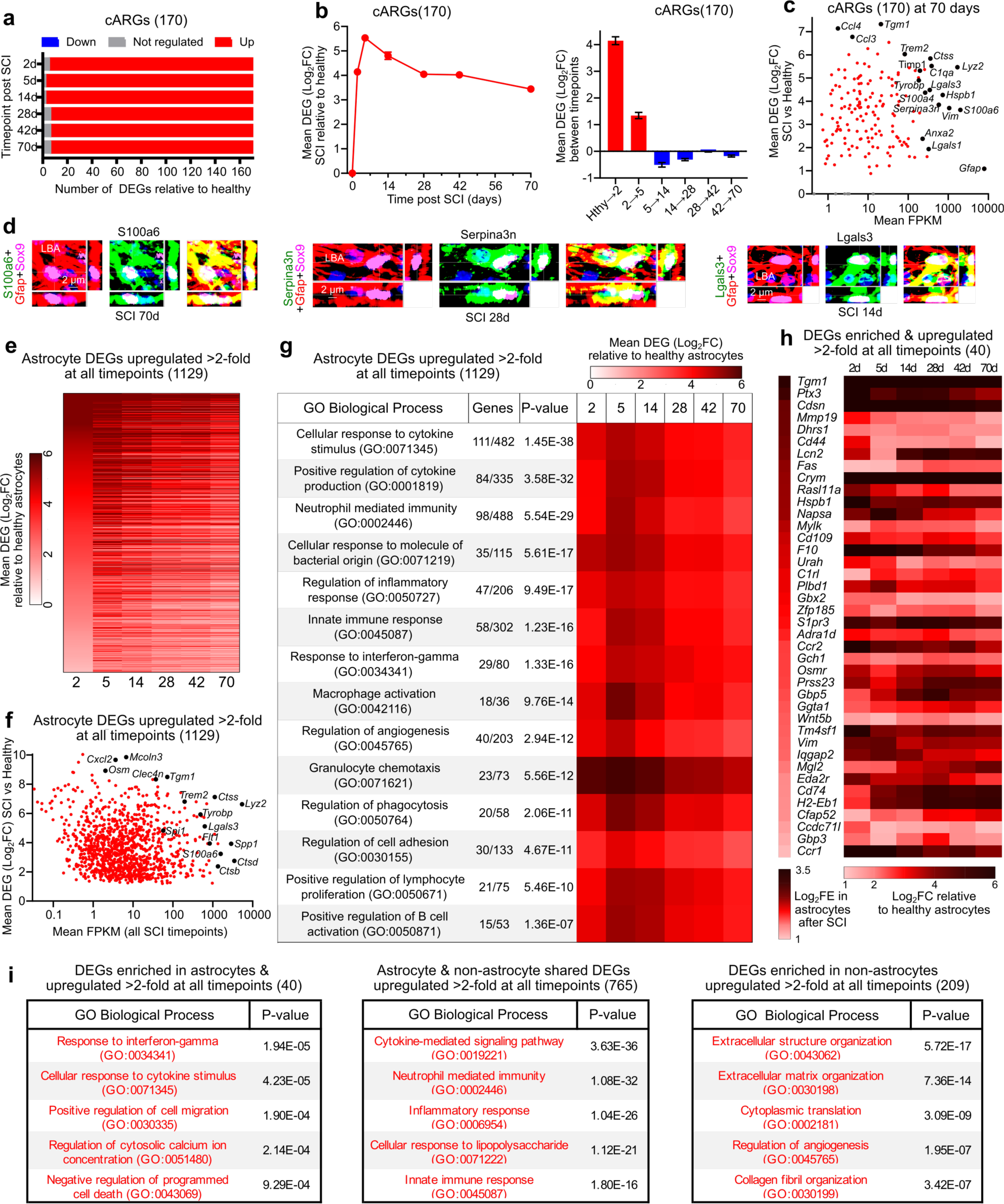
SCI-induced astrocyte transcriptional changes associated with reactivity and gains of functions. **a.** Numbers of up, down or non-significant (FDR <0.01) changes in 170 consensus astrocyte reactivity genes (cARGs) at different times after SCI. **b.** Mean log2 fold changes (log_2_FC) of upregulated cARGs at different times after SCI and changes between individual timepoints. **c.** Scatter plot showing log_2_FC and FPKM of cARGs at 70d after SCI. Selected examples of highly expressed and highly upregulated DEGs are labelled. **d.** Examples of cARGs with high levels of protein immunoreactivity in lesion border astrocytes (LBA). **e.** Heatmap of 1129 astrocyte DEGs upregulated least 2-fold at all times after SCI. **f.** Scatter plot showing log_2_FC and FPKM of 1129 astrocyte DEGs upregulated least 2-fold at all times after SCI. **g.** Top GO Biological Processes associated with 1129 astrocyte DEGs upregulated at least 2-fold at all times after SCI. **h.** Heatmaps of log2 fold enrichment (log_2_FE) and log_2_FC of 40 DEGs enriched in astrocytes by a mean of at least 2-fold compared with other cells and upregulated by at least 2-fold at all timepoints after SCI. **i.** Top GO Biological Processes associated either with 40 DEGS expressed more highly by astrocytes than other cells, or with 765 DEGs expressed at similar levels by astrocytes and other cells, or with 209 DEGs expressed more highly by other cells.

Because the injury response involves many cell types^3^, we compared expression of the same genes by astrocytes and other cells (Extended Data Fig. 3a). Many highly upregulated cARGs (Fig. 4c) were also highly enriched in reactive astrocytes compared with other cell types, such as *Tgm1*, *Steap4*, *Serpina3n* and others (Extended Data Fig. 4c). Nevertheless, many highly upregulated cARGs were de-enriched in astrocytes relative to other cells, such as *Tyrobp*, *Trem2*, *C1qc, Ccl4* and others, indicating that although these genes were used by reactive astrocytes they were also prominently used by other cells (Extended Data Fig. 4d).

We next examined 1129 DEGs upregulated by at least 2-fold at all timepoints after SCI (Fig. 4e,f). Most of these DEGs had peak expressions at 5 days and remained markedly elevated for at least 70 days (Fig. 4e). We then tracked over time the mean expression changes of astrocyte DEGs associated with representative examples of the top GO-BP associated with these 1129 DEGs (Fig. 4g). Remarkably, all top 15, and 40 of the top 50 GO-BP involved regulation of inflammation, including both innate and adaptive immune responses, such as cytokine production, granulocyte chemotaxis, macrophage activation, and lymphocyte regulation (Fig. 4g). Other persistently upregulated GO-BP included regulation of angiogenesis, phagocytosis, and cell adhesion (Fig. 4g).

To identify functions that might be preferentially associated with reactive astrocytes, we compared the GO-BP that were associated with upregulated DEGs enriched either in astrocytes or enriched in other cells. As noted above, ipAEGs exhibited a mean downregulation of DEGs, but included some upregulated DEGs (Extended Data Fig. 3b) of which only 40 were also enriched in astrocytes by over 2-fold (Fig. 4h). The top GO-BP associated with these 40 upregulated and astrocyte enriched ipAEGs included response to interferon-gamma and cytokines, cytosolic calcium regulation, and negative regulation of programmed cell death (Fig. 4i). The top GO-BP associated with DEGs that were enriched and upregulated in other cells (non-astrocytes) were reorganization of extracellular matrix (ECM) and angiogenesis, and the top GO-BP associated with DEGs similarly upregulated by both astrocytes and non-astrocytes were cytokine signaling and regulation of innate immunity and inflammation (Fig. 4i). These findings indicated that gene expression changes in astrocytes and non-astrocytes contribute to overlapping as well as to differing biological functions after SCI. Notably, ECM reorganization was more prominently associated with DEGs deriving from non-astrocytes (Fig. 4i). Chondroitin sulphate proteoglycans (CSPGs) are ECM components that have received prominent attention in SCI and have previously been attributed primarily to astrocytes. Nevertheless, three of six CSPGs were more highly expressed by non-astrocytes at all timepoints after SCI, and not a single CSPG exhibited prominent or persistent upregulation by astrocytes above baseline healthy levels (Extended Data Fig. 4e).

These findings show that as newly proliferated astrocytes reprogram after SCI, they exhibit both transient and persistent upregulation of many DEGs. Analyses of GO-BP associated with persistently upregulated DEGs points towards these reprogrammed astrocytes adopting new functional states after injury that are not prominently active or prioritized in healthy tissue, which contribute to innate immune responses, regulation of inflammation, control of infection, debris phagocytosis and regulation of angiogenesis, which we examined further.

### Immune regulation, antigen presentation and antimicrobial defense

The most prominent gain-of-function GO-BPs associated with astrocyte transcriptional reprogramming after SCI were related to innate and adaptive immune responses such as neutrophil recruitment, microglial and macrophage activation, antigen processing and presentation, anti-microbial activity, and B and T cell recruitment (Figs. 2i,3g,4g). To examine the time course of individual DEG changes related to these functions, we compiled a list of 2766 unique genes associated with these GO-BPs. Of these 2766 genes, 1708 exhibited changes by astrocytes after SCI, which consisted primarily of acute upregulation at 2 to14 days, with some returning towards baseline levels, and many exhibiting long-term upregulation for up to 70 days (Fig. 5a,b).

**Fig. 5.**
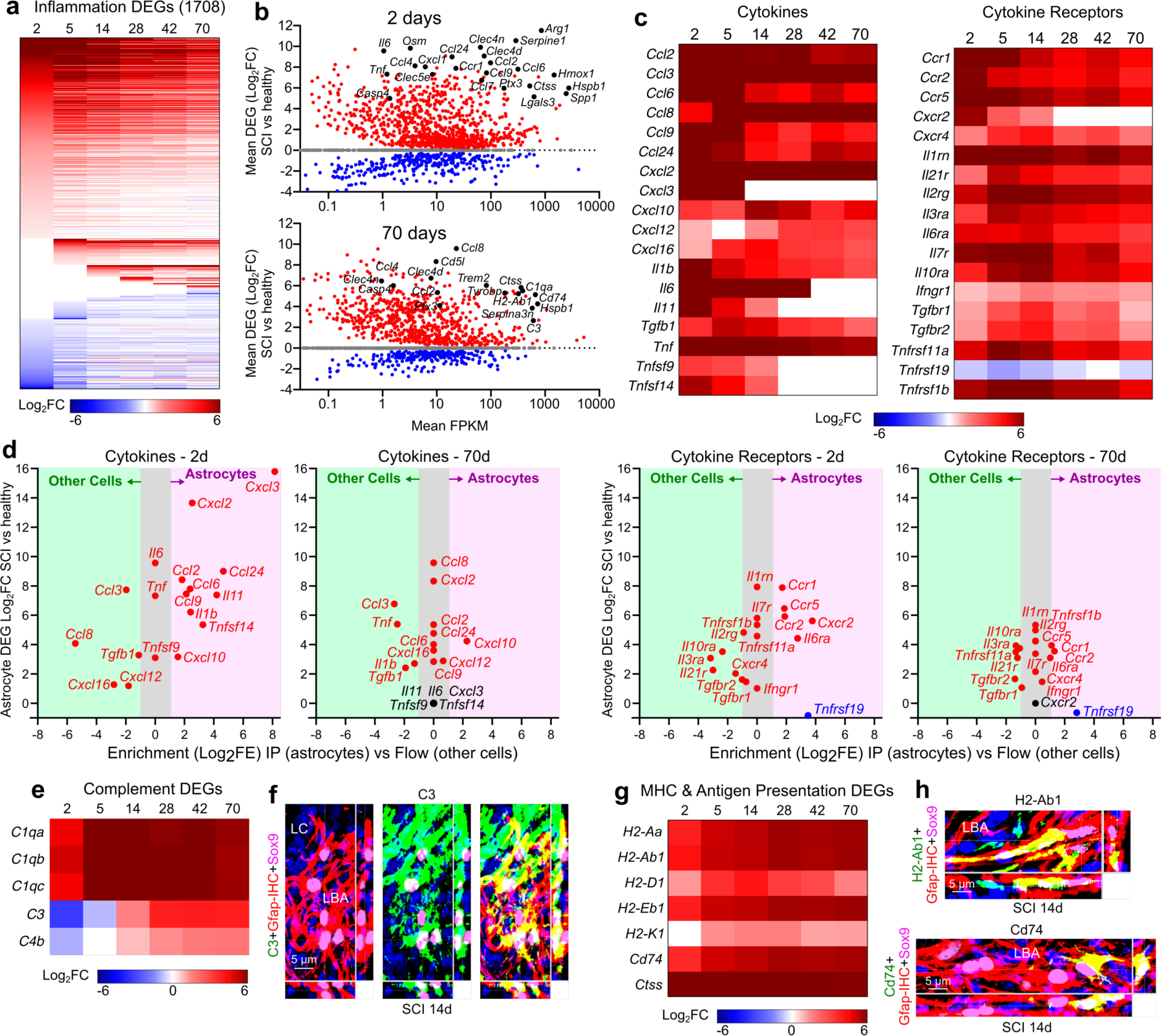
Innate and adaptive immune signaling, complement signaling and antigen presentation. **a.** Heatmap of 1708 consensus inflammation-associated DEGs up or down regulated by astrocytes at different days after SCI. **b.** Scatter plots showing log_2_FC and FPKM of 1708 astrocyte consensus inflammation-associated DEGs at 2 or 70 days after SCI. **c.** Heatmaps of mean log_2_FC of selected cytokines and cytokine receptors expressed by astrocytes after SCI. **d.** Comparison of relative enrichment and expression levels of various cytokines and cytokine receptor by astrocytes and by other cells at 2 and 70 days after SCI. **e.** Heatmap of mean log_2_FC of selected complement related DEGs expressed by astrocytes after SCI. **f.** High levels of C3 protein immunoreactivity in lesion border astrocytes (LBA) after SCI. **g.** Heatmap of mean log_2_FC of selected DEGs associated with antigen presentation and expressed by astrocytes after SCI. **h.** High levels of H2-Ab1 or Cd74 protein immunoreactivity in scattered individual LBA intermingled among many negative LBA after SCI.

In addition to multiple cytokines, chemokines and their receptors, notable immune regulatory DEGs acutely upregulated by astrocytes at 2 days after SCI included *Arg1, Ctss, Hmox1, Hspb1, Serpine1 and Lgals3;* and persistently upregulated immune regulatory DEGs at 70 days included *Cd74, H2-Ab1, H2-Eb1, H2-Aa, C1qa, C3, Trem2, Tyrobp, Serpina3n, Lcn2 and Ctss* (Fig. 5b,c). Cytokines included many pro-inflammatory, but also some anti-inflammatory (*IL11, Tgfb, IL6*), molecules (Fig. 5b,c). Most cytokines peaked rapidly followed by some decline and moderate long-term persistence (*Ccl2, Ccl3, Cxcl2, Tnf*), some cytokines peaked acutely and were downregulated to baseline (*Cxcl3, IL6, Il11*) and a few gradually increased their expression over time (*Ccl8*) (Fig. 5c). Upregulated cytokine and chemokine receptors included *Ccr1, Ccr5, 1l1rn,* and *Il10ra,* (Fig. 5c). We found evidence consistent with astrocyte pyroptosis and inflammasome generation^46^ with upregulation of *Dsdmd, Nlrp3, Pycard, Casp1, Casp 4, and Casp 8* (Extended Data Fig. 5a). In addition, multiple Toll-like receptors (Tlrs) were markedly upregulated by astrocytes from 2 to 14 days, including *Tlrs 1,4,6,7,8,9,* and some of these remained persistently elevated, (Extended Data Fig. 5b), consistent with astrocyte involvement in diverse innate immune responses.

We compared expression of the same cytokines and receptors in astrocytes and other cells (Fig. 5d). At 2 days after SCI, *Cxcl3* and *Cxcl2* were the most highly upregulated and highly enriched in astrocytes, suggesting potentially unique roles for astrocytes compared with other cells (Fig. 5d). Additional cytokines both highly upregulated and enriched in astrocytes at 2 days included *Ccl2, Ccl6, Ccl9, Ccl24, Cxcl10, Il1b, Il11, Tnfsf14* (Fig. 5d). Notably, many cytokines were upregulated by astrocytes but were nevertheless more highly expressed by other cells at 2 days, such as Ccl3, Ccl8, and Cxcl16 (Fig. 5d). By 70 days, *Cxcl3* had returned to baseline in astrocytes, whereas *Cxcl2, Ccl8, Ccl2* and *Cxcl2*, remained upregulated at levels comparable to other cells and only *Cxcl10* was persistently upregulated and enriched in astrocytes, as confirmed also by immunohistochemistry (Fig. 5d; Extended Data Fig. 5c). At 2 days after SCI, upregulated cytokine receptors enriched in astrocytes included *Ccr1, Ccr2, Ccr5, Cxcr2,* and *Il6ra,* whereas at 70 days various receptors were upregulated but were present at levels comparable to other cells (Fig. 5d). These findings suggest the potential for certain unique, and many shared, cytokine-related functions among astrocytes and other cell types after SCI.

Astrocytes also upregulated genes associated with antimicrobial defense in complement pathways and antigen presentation. *C1qa,b,c* were upregulated by over 50-fold from 5 days through at least 70 days after SCI (Fig. 5e). Microbiocidal^47^ *C3* and *C4b* were initially downregulated at 2 and 5 days but were then persistently upregulated through 70 days (Fig. 5e), and immunoreactive C3 protein was prominently detected in lesion border astrocytes that interfaced with non-neural lesion core cells in SCI and stroke (Fig. 5f; Extended Data Fig. 5d). Notably, although C1q was upregulated by astrocytes it was more highly expressed by other cells, whereas C3 was both upregulated and enriched in astrocytes (Extended Data Fig. 5e). Complement receptors *C3ar1* and *C5ar1* were also upregulated by over 50-fold and were initially enriched in astrocytes but became equally or more highly expressed by other cells by 28 days and longer after SCI (Extended Data Fig. 5e).

Additional anti-microbial defense DEGs included *Clec4d, Clec4n, Clec5a, Clec7a,* that encode pathogen-associated molecular patterns (PAMPs) receptors for viruses, bacteria and fungi, as well as *Ptx3*, *Lyz2* and *Lgals3* (Figs. 4c,d, 5b; Extended Data Figs. 5f,g). Multiple DEGs associated with antigen presentation and major histocompatibility complex (MHC) Class II were not only prominently upregulated from 5 through 70 days after SCI, but were in many cases enriched in astrocytes compared with other cells, including *H2-Aa, H2-Ab1, H2-Eb1, Cd74* (Fig. 5g,h; Extended Data Fig. 5h,i). Notably, immunoreactive protein for these MHC Class II and related molecules was high in scattered lesion border astrocytes while not detectable in others (Fig. 5h; Extended Data Fig. 5h) consistent with specialized expression among some but not other astrocytes.

These findings show that the transcriptional reprogramming of newly proliferated astrocytes after SCI includes both transient and persistent changes in many DEGs associated with multiple innate and adaptive immune functions, including diverse pro-and some anti-inflammatory cytokine signaling, antigen presentation and antimicrobial defense. The persistent upregulation of these DEGs points towards a long-term contribution to immune preparedness by border forming astrocytes around persisting lesions.

### EMT, wound repair, cell adhesion and maturation

Another prominently upregulated GO-BP associated with ipAEGs was ‘Regulation of epithelial mesenchymal transition (EMT)’ (Fig. 3f). We examined a panel of 197 consensus EMT-associated DEGs adapted from Msigdb gene sets^40^ and found a rapid acute increase in mean expression that peaked at 2 to 5 days and declined thereafter but remained persistently elevated above baseline, including prototypical EMT genes such as *Vim, Fn1, Acta2* (Fig. 6a-c). Previous studies have identified EMT genes in astrocyte responses to SCI^40,48,49^ and EMT-associated DEGs in adult cells have been implicated in wound healing^49–51^. We therefor conducted a hypothesis-driven analysis of 278 DEGs associated with the GO-BP ‘Wound healing’ (GO:0042060) and also found a rapid acute increase in mean expression in astrocytes at 2 to 5 days after SCI that declined gradually but remained persistently elevated (Fig. 6d). Notable and unexpected wound healing-associated DEGs that were highly expressed by astrocytes sub-acutely from 2-14 days after SCI included: coagulation factors *F7*, *F10, F13a1* that were upregulated over 50 to 100-fold and enriched in astrocytes compared with other cells (Fig. 6e; Extended Data Fig. 6a); heparin degrading and hemostasis molecules *Pf4, Hpse*, *Fermt*; heme degrading enzyme *Hmox1;* phagocytosis promoting *Cd44*; membrane repair *Dysf*; tissue remodeling *Mmp12*, *Timp1*; and free-radical scavenger *Gpx1;* and others including *Hbegf, Cd109, Cd151, Mylk, Pdpn* (Fig. 6e,f). Remarkably, many of these DEGs were both upregulated and enriched in astrocytes compared with other cells, suggesting unique and important wound repair functions for reactive astrocytes (Extended Data Fig. 6c).

**Fig. 6.**
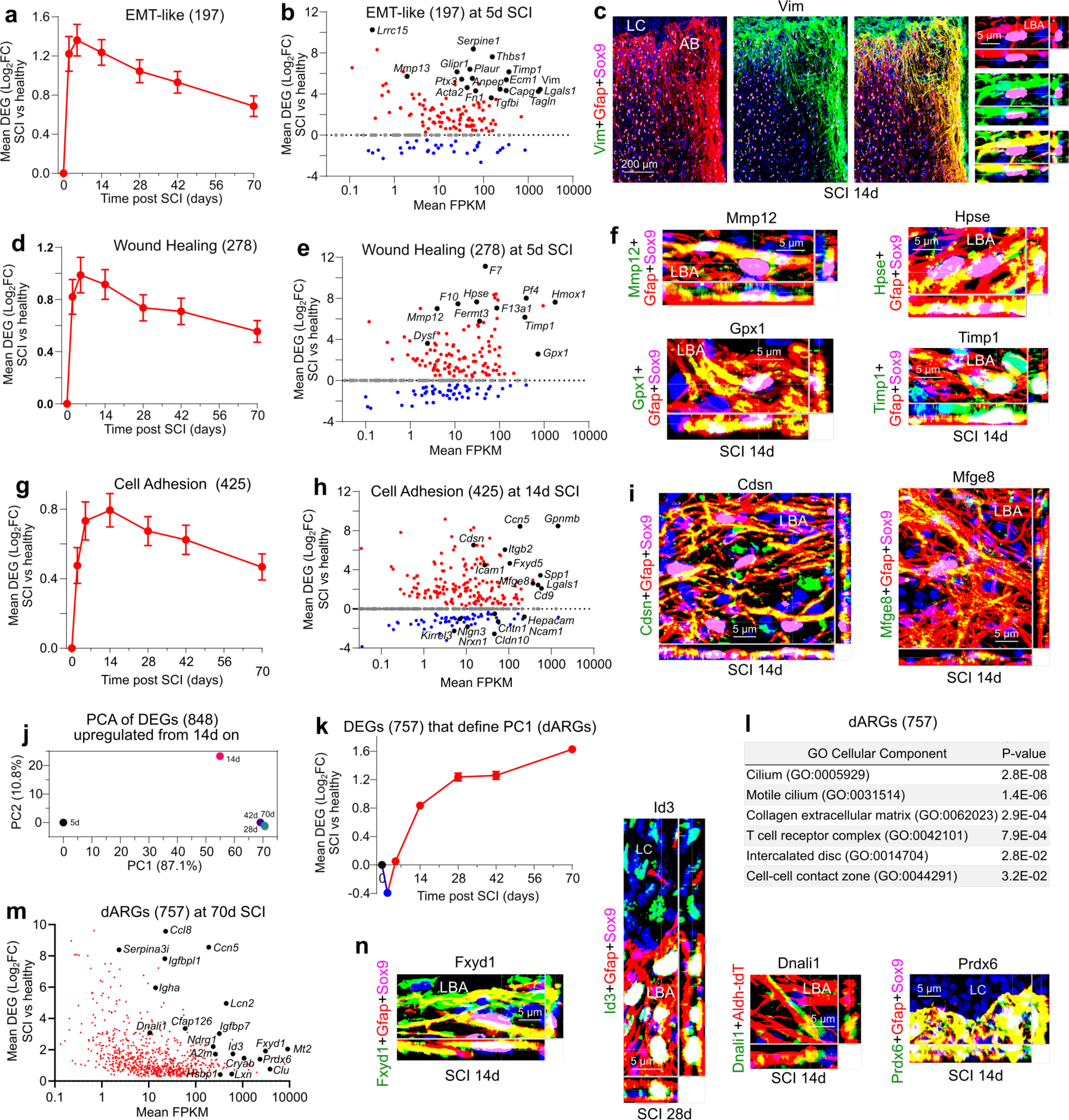
Wound healing, cell adhesion changes and border maturation. **a.** Mean log2 fold changes (log_2_FC) of a panel of 197 EMT-like DEGs at different times after SCI relative to uninjured. **b.** Scatter plot showing log_2_FC and FPKM of EMT-like DEGs at 5d after SCI. **c.** High protein immunoreactivity of the canonical EMT marker, Vim, in lesion border astrocytes (LBA). **d.** Mean log_2_FC of a panel of 278 wound healing associated DEGs at different times after SCI. **e.** Scatter plot showing log_2_FC and FPKM of wound healing DEGs at 5d after SCI. **f.** High protein immunoreactivity of selected wound healing associated molecules in LBA. **g.** Mean log_2_FC of a panel of 425 cell adhesion molecule (CAM) DEGs at different times after SCI. **h.** Scatter plot showing log_2_FC and FPKM of CAM DEGs at 14d after SCI. **i.** High protein immunoreactivity of selected CAMs in LBA. **j.** PCA of 848 DEGs upregulated from 14d onwards after SCI relative to 5d after SCI. **k.** Mean log_2_FC of the 757 delayed astrocyte reactivity genes (dARGs) that define PC1. **l.** Top GO Cellular Components associated with the 757 dARGs. **m.** Scatter plot showing log_2_FC and FPKM of dARGs at 70d after SCI. **n.** Examples of high protein immunoreactivity in LBA of selected dARGs.

As noted above, newly proliferated lesion border astrocytes downregulate the domain-associated cell adhesion molecule (CAM), *Hepacam*^42^, (Fig. 3d) and as they mature do not adopt individual domains but instead reorganize with highly overlapping and intermingled cell processes (Fig. 1). We therefor looked for changes in DEGs associated with cell-cell interactions by examining a panel of 425 CAMs and cell-cell interaction associated genes compiled from GSEA, GO and literature. We found a pronounced early and persisting overall mean upregulation in expression of CAM-associated DEGs, but also some down regulation by astrocytes after SCI (Fig. 6g-i; Extended Data Fig. 7a). Notable upregulated CAM-associated DEGs included, *Ccn5, Gpnmb, Spp1* involved in integrin binding; *Lgals1, Fxyd5* involved in matrix adhesion; *Cdsn, Cd9, Mfge8* involved in cell-cell adhesion; and *Itgb2, Icam1* involved in leukocyte adhesion (Fig. 6h,i; Extended Data Fig. 7b,c). Notable downregulated CAMs included *Hepacam, Cldn10, Cntn1, Kirrel3, Ncam1, Nrxn1, Nlgn3* involved in homophilic astrocyte-astrocyte interactions and interactions with neurons (Fig. 6h). Many of these CAMs were enriched in astrocytes compared with other cells (Extended Data Fig. 7d).

The DEGs and associated GO-BPs examined thus far, such as immune regulation and wound healing, peaked acutely after SCI. To identify potential features of border forming astrocytes that might emerge after the acute period, we next examined 848 DEGs that were upregulated from 14 to 70 days. PCA revealed a prominent difference defined by PC1 between 5 days and the more chronic timepoints, and smaller difference defined by PC2 between 14 days and 28 to 70 days (Fig. 6j). Factor analysis of the 757 DEGs defining PC1 revealed a cohort of delayed astrocyte reactivity genes (dARGs) whose mean expression declined acutely and then increased and remained high from 14 to 70 days (Fig. 6k). These dARGs exhibited GO Cellular Components associated with ciliated cells, extracellular matrix interactions, and intercalated disk cell-cell contact interactions (Fig. 6l). Notable dARGs at 70d after SCI included the transcription factor *Id3*; proteinase inhibitors *A2m, Lxn, Serpina3i*; antioxidants *Mt2, Prdx6*; ion transport regulator *Fxyd1*; chaperone *Clu*, multifunctional *Igfbpl1*, antimicrobial *Lcn2, Nrdg1, Igha*; heat shock protein *Cryab*; cilia associated proteins including *Dnali1, Cfap126* (Fig. 6m,n; Extended Data Fig. 7e,f). Many of these dARGs were enriched in astrocytes (Extended Data Fig. 7d), and protein expression for various examples was confirmed by immunohistochemistry (Fig. 6n; Extended Data Fig. 7e). Delayed activation of *Id3* and multiple cilia related genes including *Dnali1, Cfap126* (Fig. 6m; Extended Data Fig. 7f) aligns with their established roles in proliferation arrest and promotion of astrocyte differentiation of immature progenitor-like cells and points towards their possible involvement in ending the transient phase of proliferation and immaturity observed in the initial post-injury response^14,52–54^.

Together these findings show that the transcriptional reprogramming of newly proliferated astrocytes after SCI equips astrocytes with the capacity to contribute to important wound healing processes, including contributions to hemostasis, coagulation, and breakdown of heme in hemorrhagic CNS wounds, as well as free-radical scavenging and tissue and matrix remodeling. Remarkably, many of the associated DEGs were not only upregulated but were also highly enriched in astrocytes compared with other cells, suggesting unique and important wound repair functions for newly proliferated reactive astrocytes. In addition, the findings reveal major changes in CAM expressions (both up and down) by lesion border astrocytes around CNS lesions, indicating major readjustments in their homophilic and heterophilic cell-cell interactions consistent with their major change in morphology from cells with unique non-overlapping cellular domains^42^ to cells with highly overlapping and intertwined cellular processes (Fig. 1c).

### Phenotypic features of mature lesion border astrocytes

We next conducted snRNAseq and immunohistochemical protein analyses to look for phenotypic features of mature lesion border astrocytes compared with astrocytes in healthy spinal cord. We isolated and sequenced 139,981 nuclei from the same region of thoracic spinal cord in uninjured mice and at 28d after SCI when borders are mature (Fig. 7a). Nuclei were filtered and annotated using panels of multiple cell-type specific marker genes, and clusters identified as astrocytes were extracted for further processing (Extended Data Fig. 8a,b). Re-clustering and further extraction based on multiple astrocyte specific marker genes confidently identified 15,637 astrocyte nuclei that were used for final analyses (Fig. 7a-d, Extended Data Fig. 8c).

**Fig. 7.**
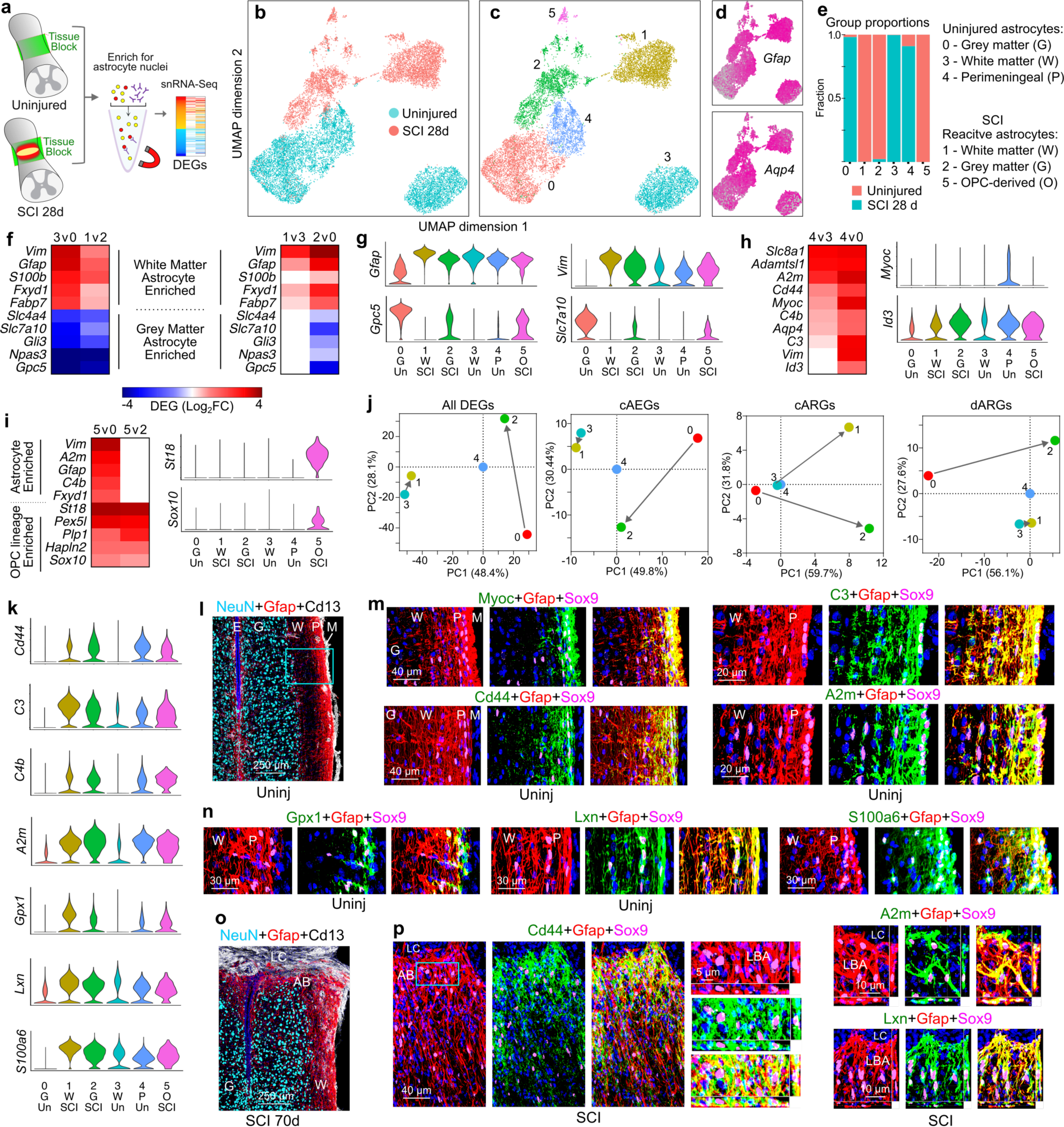
Shared and distinct molecular features of uninjured astrocytes and mature lesion border astrocytes (LBA). **a.** snRNAseq procedures. **b,c.** UMAP clusters of astrocyte nuclei. **d.** *Gfap* and *Aqp4* expression across clusters. **e.** Proportion of nuclei from uninjured (Un) or SCI mice per cluster. **f,g.** Heatmaps (f) and violin plots (g) comparing marker genes enriched in grey or white matter astrocytes across different clusters. **h,i.** Heatmaps and violin plots of genes enriched in perimeningeal astrocytes (h) or in OPC-derived astrocytes. **j.** PCA comparing changes in different gene cohorts across clusters after SCI. **k.** Violin plots of genes enriched in both perimeningeal and reactive astrocytes compared with uninjured. **l-n.** Immunohistochemistry of proteins enriched in uninjured perimeningeal astrocytes (P) and LBA after SCI. **l,o.** Survey images of uninjured (l) and SCI (o). **m,n.** Details from boxed region in l showing proteins enriched in perimeningeal astrocytes. **p.** Proteins enriched in LBA. E, ependyma, LC, lesion core, M, meninges.

UMAP analysis showed pronounced separation of uninjured and SCI derived nuclei, and we categorized six major clusters of astrocytes, three from uninjured and three from SCI (Fig. 7b,c). Canonical markers of healthy and reactive astrocytes such as *Gfap* and *Aqp4* were expressed throughout all clusters (Fig. 7d). Consensus markers discriminated healthy grey (cluster 0) and white (cluster 3) matter astrocytes and identified separate major clusters of reactive astrocytes with transcriptional features associated with grey (cluster 2) and white (cluster 1) matter astrocytes (Fig. 7e-g). Healthy spinal cord astrocytes also separated into a third cluster (cluster 4) with selective or enriched expression of molecules such as *Myoc, Cidea, and Id3* (Fig. 7h; Extended Data Fig. 9a) that have previously been associated with perimeningeal astrocytes that form so-called *’limitans’* borders along the surface of the entire CNS^40,55^. Immunohistochemistry confirmed robust protein expression of Myoc and Id3 proteins in healthy perimeningeal astrocytes (Fig. 7m; Extended Data Fig. 9b).

An additional small cluster (cluster 5) selectively expressed both reactive astrocyte markers and OPC lineage markers *St18, Sox10, Plp1, Hapln2, Ninj2, Enpp6* (Fig. 7g,i; Extended Data Fig. 9c). Thus, our findings confirm by both snRNAseq and lineage tracing (Fig. 1; Extended Data Fig. 1) the interesting observation^32–35^ that local OPC can give rise to a small contingent of lesion border astrocytes, noted also in another recent single cell RNAseq study^56^.

We next evaluated how different gene cohorts consisting of all DEGs, cAEGs, cARGs or dARGs changed in the different major astrocyte clusters after SCI. PCA analyses showed that in all cases, grey matter astrocytes exhibited the greatest changes at 28 days after SCI including markedly reduced expression of cAEGs, and markedly upregulated cARGs and dARGs (Fig. 7j). In contrast, white matter astrocytes exhibited minimal changes in gene expressions except for markedly upregulating cARGs (Fig. 7j). These findings suggest that the persistent overall downregulation of cAEGs noted above (Fig. 4b-e) is associated more with grey matter than white matter astrocytes. Interestingly, with respect to most gene cohorts, healthy perimeningeal astrocytes (cluster 4) were closer in PC space to reactive astrocytes than to healthy grey or white matter astrocytes. Nevertheless, when focusing specifically on cARGs, perimeningeal astrocytes grouped much closer to healthy astrocytes (Fig. 7j).

Because various lines of evidence have historically suggested similarities between reactive astrocytes that form borders around CNS lesions and *’limitans’* astrocytes that form borders with fibroblast lineage meningeal cells along the entire CNS surface^4,57–59^, we looked for potential similarities and differences in gene expression and protein immunoreactivity. Notable molecules enriched in both healthy perimeningeal astrocytes (cluster 4) and lesion border reactive astrocytes (clusters 1,2,5) compared with healthy grey or white matter astrocytes included *Id3*, *Cd44, C3, C4b, A2m, Gpx1, Lxn, Vim, S100a6, Lgals3, Fxyd1, Padi2, Prdx6* (Figs. 7g,h,k-p; Extended Data Fig. 9b,d-f). Both *Myoc* and *Cidea* were selective for healthy perimeningeal astrocytes and were essentially not detected in reactive astrocytes or in healthy grey or white matter astrocytes (Figs. 7h,m; Extended Data Fig. 9a). Conversely, healthy perimeningeal astrocytes did not express multiple markers associated with overt astrocyte reactivity, including *Lyz2, Serpina3n, Fcerg1, Ctss, Trem2, Tyrobp, C1qa, Hsbp1* and others (Fig. 8a, Extended Data Figs. 8c, 9g). Reactive perimeningeal astrocytes were not discriminated as a separate cluster most likely because they become indistinguishable from other reactive astrocytes due to the pronounced and persistent downregulation of *Myoc* and other genes that distinguish them (Extended Data Fig. 9h). These findings indicate that healthy perimeningeal astrocytes exhibit unique features that distinguish them from healthy grey and white matter astrocytes and share some features with reactive astrocytes but do not exhibit overt reactivity profiles.

**Fig. 8.**
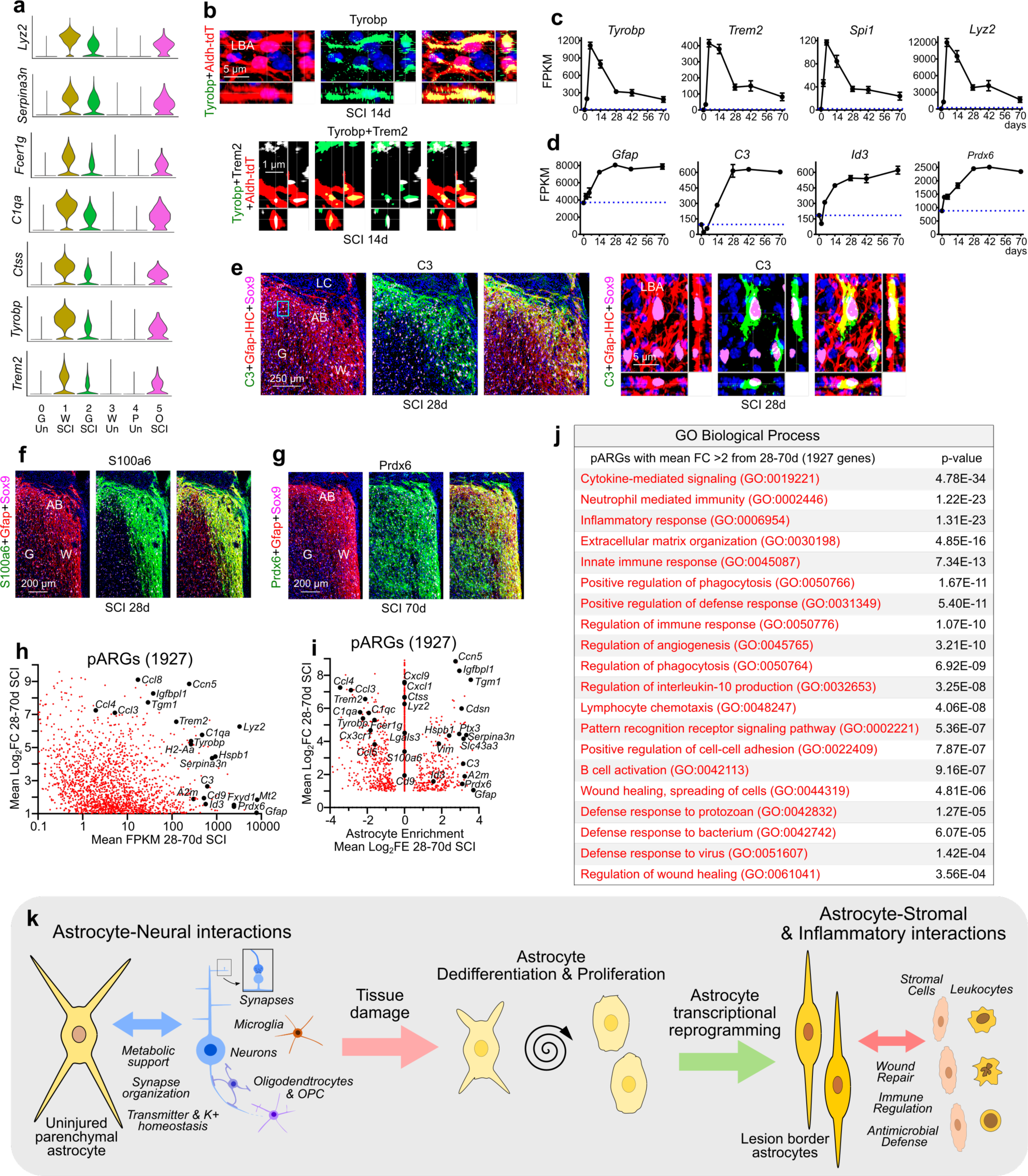
Features of mature lesion border astrocytes (LBA). **a.** Violin plots of snRNAseq detected genes enriched in reactive astrocyte clusters compared with uninjured. **b.** Immunohistochemistry of proteins Tyrobp and Trem2 in LBA after SCI. **c,d.** Selected astrocyte DEGs exhibiting different patterns of expression changes in the form of acute rise followed by decline (c) or delayed but persistent increase (d) after SCI as detected by Astro-RiboTag RNAseq. **e,f,g.** Immunohistochemistry of proteins C3, S100a6 and Prdx6 in LBA after SCI. Box in e shows location of detail. **h.** Scatter plot comparing mean log_2_FC and mean FPKM of 1927 pARGs upregulated at least 2-fold from 28-70d after SCI as detected by Astro-RiboTag RNAseq. **i.** Scatter plot comparing mean log_2_FC and mean enrichment (log_2_FE) of 1927 pARGs upregulated from 28-70d after SCI. **j.** GO Biological Processes associated with 1927 pARGs upregulated at least 2-fold from 28-70d after SCI. **k.** Summary schematic showing local astrocyte responses to CNS tissue damage by dedifferentiation, proliferation and transcriptional reprogramming into border forming wound repair astrocytes.

To further examine phenotypic features of mature lesion border reactive astrocytes, we examined more closely timepoints from 28 to 70 days after SCI from our Astro-RiboTag data set (Fig. 8c-j). We noted two common but different gene expression patterns. Certain reactivity DEGs peaked acutely from 2 to14 days and then declined by 28 days but continued to be expressed thereafter at levels higher than in healthy astrocytes, such as *Tyrobp, Trem2, Spi1, Lyz2, Lgals3, Ccl4* and others (Fig. 8c; Extended Data Fig. 9i), and DEGs with this pattern were generally not shared with perimeningeal astrocytes (Fig. 8a, Extended Data Fig. 9g). In contrast, certain other reactivity DEGs only reached their peak expression from 28 to 70 days and then persisted at those levels, such as *Gfap, C3, Id3, S100a6, Prdx6* and others (Fig. 8d; Extended Data Fig. 9i). Notably, this pattern was particularly exhibited by DEGs shared with perimeningeal astrocytes that also form borders to stromal cells (Figs. 7k-p; Extended Data Fig. 9b,d-f,j). Notably, certain transcriptional regulators mirrored these changes and with an early peak and then decline in expression, such as *Spi1* (Fig. 8c), while other transcriptional regulators exhibited delayed and persisting increases in expression, such as *Id3* (Fig. 8d).

Although *Trem2* and *Tyrobp* were not detectably expressed in healthy astrocytes they were robustly upregulated by reactive astrocytes after SCI as detected by both snRNAseq and Astro-RiboTag RNAseq, and we confirmed protein expression by immunohistochemistry (Fig. 8a-c,h; Extended Data Fig. 8c). We have previously shown that reactive astrocytes can adopt gene expressions that are unexpected based on absence of expression in healthy CNS, such as the transcription factor *Spi1*^16^ (Fig. 8c). *Trem2, Tyrobp* and *Spi1* are widely regarded as uniquely expressed by microglia in the CNS^60,61^, but can be expressed by reactive astrocytes albeit at levels lower than other cells (Fig. 8i).

The antimicrobial defense factor *C3*^62^, became persistently upregulated by over 6-fold from 28 to 70 in astrocytes after SCI and immunoreactive protein was highly concentrated in lesion border astrocytes that separate lesion core tissue from immediately adjacent spared grey matter (Fig. 8d,e). C3 has been proposed as an unequivocal marker of astrocytes that are neurotoxic to neurons in CNS disorders. However, *C3* is expressed by certain astrocytes in healthy tissue, particularly perimeningeal astrocytes, and then declines acutely after SCI during periods when neurons are lost, after which *C3* becomes highly and persistently upregulated in astrocytes that separate inflamed tissue lesions from immediately adjacent surviving neurons (Figs. 7o, 8d,e). These findings suggest that *C3* is expressed by reactive astrocytes to exert natural antimicrobial functions^62^ at sites of tissue damage, and challenge the proposal that *C3* is an unequivocal marker of rogue ‘neurotoxic’ astrocytes.

Certain DEGs such as the calcium binding protein *S100a6*, which are not expressed by healthy grey matter astrocytes became highly upregulated by lesion border astrocytes that exhibit grey matter features (Cluster 2) and immunoreactive S100a6 was prominent in astrocyte borders along lesions adjacent to spared grey matter (Figs. 7k,8f; Extended Data Fig. 9i). Other DEGs such as the antioxidant *Prdx6,* and the intercalated disc protein *Fxyd1* that were expressed by healthy grey and white matter astrocytes as well as perimeningeal astrocytes, became prominently and persistently upregulated by lesion border astrocytes (Figs. 8d,g; Extended Data Fig. 9e,f,i,j). Notably, lesion border astrocytes and perimeningeal astrocytes shared expression of multiple immunoreactive proteins such as Cd44, A2m, Id3, C3, S100a6, and Gpx (Fig. 7l-p), some of which were particularly enriched in border-forming astrocytes and expressed at low or undetectable levels in healthy astrocytes and have the potential to serve as markers of border astrocytes (Extended Data Fig. 10a,b).

To identify phenotypic features that might be preferentially associated with mature reactive lesion border astrocytes compared with other cells in the lesion area, we examined a cohort of 1927 DEGs that were persistently up regulated by greater than 2-fold from 28 to 70 days, which we refer as persisting astrocyte reactivity genes (pARGs) (Fig. 8h). We then determined the enrichment of pARGs in either astrocytes or in other cells (Fig. 8i). Interestingly, the majority of pARGs that were highly upregulated by astrocytes were either expressed at equal (1330 pARGs or greater (335 pARGs) levels by other cells, including *Tyrobp, Trem2, Lyz2, Lgals3, Cxcl1, Cxcl9, Ccl3, Ccl4, C1qa, C1qc, Fecer1g* and others, and most of these pARGs had peak expression levels acutely after SCI and then declined but continued to be persistently expressed by mature lesion border astrocytes (Fig. 8i) (Fig. 8c,i; Extended Data Fig. 9i). Nevertheless, 262 pARGs were both highly upregulated and highly enriched in astrocytes including, *Gfap, Tgm1, Ptx3, A2m, C3, Cdsn, Serpina3n, Hspb1, Id3, Timp1, Vim* and others, and most of these pARGs had higher expression levels chronically than acutely after SCI (Fig. 8c,i; Extended Data Fig. 9j). These findings indicate that gene expression changes in astrocytes and other cells are likely to contribute to various overlapping functions after SCI. For example, many of the pARGs upregulated by astrocytes but expressed more highly by other cells were involved in innate immune responses and regulation of inflammation (Fig. 8h,i). Thus, reactive astrocytes contributed multiple molecules to the regulation of inflammation but were not necessarily the main producers of those molecules (Fig. 8i). Lastly, we examined the top GO-BP associated with all 1927 pARGs, which prominently included multiple functions associated with innate immune responses, regulation of inflammation, wound healing, and anti-microbial defense (Fig. 8j).

## Discussion

In this study, we show that after CNS injury, local mature astrocytes dedifferentiate, proliferate, and become transcriptionally reprogrammed into persisting new functional states. They down-regulate molecules associated with homeostatic astrocyte-neuron interactions, and upregulate molecules associated with wound healing, immune regulation, and microbial defense (Fig. 8k). Thus, these locally derived, newly proliferated, and reprogrammed astrocytes adopt canonical features of essential wound repair cells that persist in adaptive states and are the predominant source of neuroprotective borders that re-establish CNS integrity by separating neural parenchyma from stromal and immune cells as occurs throughout the healthy CNS.

Notably, astrocyte transcriptional profiles changed markedly over time after injury such that certain DEGs exhibited maximal differences early after injury and then returned towards, but not quite reaching, baseline healthy levels, while other DEGs exhibited delayed and persistent increases indicative of permanent reprogramming to new astrocyte functional states. Expression levels of certain transcriptional regulators mirrored these changes respectively, such as *Spi1* that peaked early and declined, and *Id3* that peaked late and persisted, in a manner consistent with context specific regulation of different astrocyte reactivity states^16^. These findings have implications for understanding the diversity of astrocyte responses to injury and disease and how this diversity is regulated in different contexts. For example, there is increasing evidence that astrocytes for unknown reasons attenuate certain homeostatic functions in aging and related neurodegenerative disorders. Our findings suggest that astrocyte downregulation of homeostatic functions may be controlled by signaling mechanisms that are naturally triggered by tissue damage as part of the dedifferentiation, proliferation, and reprogramming that is fundamental to mounting a wound repair response. In this regard our findings suggest potential links of such responses to neurodegenerative conditions such as Alzheimer’s disease where astrocyte loss- and gain-of-functions may contribute to disease progression^63^, and where the top 30 most consistently upregulated proteins across multiple studies included at least 11 molecules identified here not only as upregulated by, but also enriched in, wound repair astrocytes: Cd44, S100a6, Padi2, Prdx6, C3, Gpx1, Hsbp1, Gpnmb, Clu, Vim, Gfap^64^. Understanding the regulation of astrocyte specific responses in different contexts will help identify ways to selectively attenuate detrimental, or augment beneficial, astrocyte reactivity responses as appropriate to improve outcomes. In this regard, we posit that astrocyte proliferation and reprogramming is probably underestimated in neuropathology because it has been difficult to reliably identify proliferated astrocytes and distinguish them from cells that have merely upregulated GFAP. The identification here of additional protein markers associated with these cells may facilitate future neuropathological studies.

We show by lineage tracing and transcriptional profiling that the vast majority of over 90% of newly proliferated border-forming astrocytes around lesions derive from local Aldh1l1-expressing astrocytes. We also confirm by lineage tracing and snRNAseq that local OPC can also give rise to GFAP-expressing lesion border astrocytes around CNS lesions^32–35^, and we show that these comprise a small proportion of about 10% of lesion border astrocytes after SCI or stroke. OPC-derived border-forming astrocytes appear morphologically similar to astrocyte-derived ones and express many molecular markers of astrocyte reactivity, but also exhibit some transcriptional differences. Future studies using techniques such as RABIDseq^65^ will be useful to examine whether OPC- or astrocyte-derived border forming cells contribute overlapping or differing functions. Ependymal cells have also been proposed as a major source of lesion border astrocytes^66^, but this could not be confirmed by previous rigorously characterized lineage-tracing, which showed that no border-forming astrocytes derive from ependymal cells after SCI^30^ or forebrain stroke^67^, and our lineage tracing here accounts for essentially all lesion border astrocytes as derived from local astrocytes or OPC.

The question arises as to why newly proliferated astrocytes form borders around CNS lesions as part of the wound repair process. Astrocyte proliferation and formation of borders around CNS lesions has long been characterized as scar formation that contributes to inferior wound repair and regeneration failure. Accordingly, preventing or removing these borders was long regarded as a major goal to improve outcome. Challenging this notion, a large and steadily accumulating body of transgenic loss-of-function studies indicates that preventing, attenuating or disrupting astrocyte lesion borders causes detrimental effects to neural wound repair, including increased spread of inflammation, persistent blood-brain barrier permeability, increased loss of neural tissue, decreased functional recovery, and in some cases increased mortality^4,16,17,22,23,25–28^, whereas augmenting border formation reduces lesion size^22^. In healthy CNS, all neural parenchyma is segregated from non-neural stromal cells either by astrocyte endfeet along blood vessels or by so-called *’limitans’* astrocytes that abut meningeal cells around the entire CNS. Structural similarities between these perimeningeal astrocytes and astrocyte borders around lesions have been noted previously^4,57–59^. Here, we extend these observations by showing that lesion-border astrocytes share molecular similarities with healthy perimeningeal astrocytes but also exhibit additional functions related to wound repair and heightened levels of microbial defense and immune regulation. The idea that astrocytes surrounding CNS lesions form new *limitans* borders rather than fibrotic scar tissue has been considered before, particularly in human pathology^57–59^. Notably, scar tissue derives from proliferating stromal cells that rapidly replace damaged organ parenchyma with connective tissue to provide structural integrity to the wound site and prevent infection, but scar tissue is unable to reconstitute lost CNS functions. In pro-regenerative mammalian organs, such as liver, intestinal epithelia and skin epidermis, injury induces parenchymal cell proliferation that enables tissue regeneration and recovery of organ function^68–70^. In contrast to CNS, proliferating parenchymal cells in these other organs are not considered to form scars; and stromal cell scarring occurs when parenchymal regeneration is inadequate^68–70^. Astrocytes are key neural parenchymal cells that derive from the same neural stem cells as neurons^7^, but unlike mature neurons and oligodendrocytes which are post-mitotic, mature astrocytes can re-enter the cell cycle and proliferate after injury to generate new astrocytes that isolate true stromal cell scar tissue and thereby re-establish neural parenchymal integrity and preserve neural tissue and function by separating neural parenchyma from stromal and immune cells as occurs throughout the healthy CNS. These findings advocate that astrocyte borders around lesions should no longer be referred to as scars.

The appealing notion of potentially achieving scar free wound healing by parenchymal cells in the adult CNS is supported by wound repair observations in mammalian neonates, where microglia and astrocytes sustain high levels of proliferation that enable a rapid glia-based repair rather than fibrotic repair, which is sufficient to promote neural circuit regeneration and recovery of function that is sustained into adulthood^71,72^. This may occur in part because neonatal astrocytes are proliferative at the time of injury and may rapidly increase proliferation to replace lost neural parenchyma without stromal scarring in a manner similar to pro-regenerative organs. In contrast, astrocytes in mature CNS are dormant with respect to proliferation, and their proliferative response to injury is delayed, spatially restricted and transient, and is adequate to generate new astrocyte borders around the rapidly formed stromal cell scars, but is insufficient to support rapid wider spread parenchymal repair. These observations support the pursuit of strategies that accelerate and extend astrocyte post-injury proliferative and immaturity states to augment neural parenchymal repair^22^ in a variety of contexts associated with neural parenchymal loss.

## Methods

### Animals

Young adult male and female C57BL/6 mice between two and four months of age at the time of experimental procedure were used for all studies. For lineage tracing, Ai14 mice expressing the reporter, tdTomato (tdT) (JAX: 007914) were crossed with different Cre-driver lines: (1) Aldh1l1-CreERT2^36^ (JAX: 031008), (2) Pdgfra-CreERT^37^ (JAX: 018280) or (3) NG2-CreERT^38,39^ (JAX: 008538). For sequencing of astrocyte ribosome associated RNA, mice expressing RiboTag^31^ (JAX: 029977) were crossed either with Aldh1l1-CreERT2^36^ (JAX: 031008) or with mGfap-Cre-73.12^73^ (JAX: 012886). For postnatal astrocyte evaluations, mGfap-RiboTag mice were used at postnatal days P0, P3, P7, P14, P21, P35, P63. Transgene expression for each sample was confirmed by genotyping of collected tail samples prior to processing for astrocyte specific RNA. Mice were housed in a specific pathogen-free facility with 12-h light/dark cycle and controlled temperature and humidity and were allowed free access to food and water. All in vivo experiments involving the use of mice were conducted according to protocols approved by the Animal Research Committee (ARC) of the Office for Protection of Research Subjects at University of California Los Angeles (UCLA). ARC Numbers: ARC-2017-044; ARC-2008-051; ARC # 2015-073; ARC-2000-001.

### Surgical procedures

All surgeries were performed under general anesthesia with isoflurane in oxygen-enriched air using an operating microscope (Zeiss, Oberkochen, Germany), and rodent stereotaxic apparatus (David Kopf, Tujunga, CA). All animals received the opiate analgesic, buprenorphine (0.1mg/kg), subcutaneously before surgery and every 12 h for at least 48 h post-surgery. Animals were coded numerically and evaluated thereafter blind to experimental condition.

#### Spinal cord injury (SCI)

Laminectomy of a single vertebra was performed at cord level T10. A timed (5 second) lateral compression, complete crush SCI was made using No. 5 Dumont forceps (Fine Science Tools, Foster City, CA) with a tip width of 0.5 mm. Daily bladder expression was performed for the duration of experiments or until voluntary voiding returned.

#### Stroke

After a small craniotomy over the left coronal suture, 1.5 μL of L-NIO (N5-(1-Iminoethyl)-L-ornithine) (Cat. No. 0546, Tocris solution) (27.4 mg/ml in sterile PBS) was injected into the caudate putamen nucleus at 0.15 μL/min using target coordinates relative to Bregma: +0.5 mm A/P, +2.5 mm L/M and −3.0 mm D/V by using a glass micropipette.

### Lineage tracing

Cell lineage-tracing was conducted by using tdT reporter protein targeted to specific cells and temporally regulated via CreERT-loxP in transgenic mice^29^. tdT expression was activated in young adult mice by administering tamoxifen (Sigma, T5648-1G, 20 mg/ml in corn oil) by intraperitoneal injection (100mg/kg, once a day) for five days followed by clearance for three weeks before SCI or stroke, so that no residual tamoxifen remained. Using this approach, tdT becomes constitutively expressed by cells in which it has been activated by Cre during the period of tamoxifen delivery, and this expression is passed on to all progeny cells. Once tamoxifen is no longer administered and has cleared, only the originally targeted cells and their progeny express tdT.

### BrdU

Bromodeoxyuridine (BrdU, Sigma), 100 mg/kg/day dissolved in saline plus 0.007N NaOH, was administered as single daily intraperitoneal injections on days 2 through 7 after SCI.

### Histology and immunohistochemistry

After terminal anesthesia by barbiturate overdose, mice were perfused transcardially with 4% paraformaldehyde (Electron Microscopy Sciences, Hatfield, PA). Spinal cords were removed, post-fixed overnight, and cryoprotected in buffered 30% sucrose for 48 hours. Frozen sections of the spinal cord were prepared in the horizontal plane at 30 μm thickness using a cryostat microtome (Leica) and processed for immunofluorescence as described^26^.

*Primary antibodies* goat anti-A2m (1:300, AF1938; R&D Systems); rabbit anti-Aldh1l1 (1:1000; Ab87117; Abcam, Cambridge, MA, USA); sheep anti-BrdU (1:800, NB-500-235; Novus); rat anti-C3 (1:400, NB200-540; Novus); goat anti-CD13 (1:600, AF2335; R&D Systems, USA); rat anti-Cd44 (1:400, 14-0441-82; Invitrogen); rat anti-CD68 (1:1000, MCA1957; Biorad, USA); rabbit anti-Cd74 (1:200, A13958; Abclonal); rabbit anti-Cdsn (1:800,13184-1-AP; Proteintech); goat anti-Cxcl10 (1:200, AF-466; Novus); rabbit anti-Dnali1 (1:500, 17601-1-AP, Proteintech); rabbit anti-Fxyd1 (1:800, A15082, Abclonal); rabbit anti-GFAP (1:2,000; GA524; Z033401-2; Dako/Agilent Tech., CA); rat anti-GFAP (1:1,000, 13-0300; Thermofisher, USA); rabbit anti-hemagglutinin (HA) (1:1000, H6908, Sigma); goat anti-Gpc5 (1:200, AF2607; R&D Systems); rabbit anti-Gpx1 (1:200, 29329-1-AP; Proteintech); goat anti-HA (1:800, NB600-362, Novus Biologicals); rabbit anti-H2-Ab1 (1:200, A18658; Abclonal); rabbit anti-Hpse (1:200,24529-1-AP; Proteintech);guinea pig anti-Iba1 (1:1000, 234004; Synaptic Systems, USA); rabbit anti-Iba-1 (1:800, 019-19741; Wako, Osaka, Japan); rabbit anti-Id3 (1:500; 9837; Cell Signaling); rabbit anti-Kcnj10 (Kir4.1) (1:400, APC-035; Alomone labs)rat anti-Lgals3 (1:200, 14-5301-82; ThermoFisher); rabbit anti-Lxn (1:500, 13056-1-AP; Proteintech); rabbit anti-Mfge8 (1:200, A12322; Abclonal); rabbit anti-Mmp12 (1:200, 22989-1-AP; Proteintech); goat anti-Myoc (1:400, AF2537; Novus); guinea pig anti-NeuN (1:1000, 266004; Synaptic Systems); rabbit anti NeuN (1:1000, ab177487, Abcam); guinea pig anti-Olig2 (1:800, ABE1024; Millipore); rabbit anti-Olig2(1:200, AB9610; Millipore); rabbit anti-Padi2 (1:300,12110-1-AP; Proteintech); rabbit anti-Prdx6 (1:500, 13585-1-AP; Proteintech); sheep anti-S100a6 (1:300, AF4584; R&D Systems); rabbit anti-S100a6 (1:200, A3461; Abclonal); goat anti-Serpina3n (1:200, AF4709; R&D Systems); goat anti-Sox9 (1:800, AF3075; R&D Systems); rabbit anti-Sox9 (1:800, 702016; ThermoFisher); goat anti-Sox10 (1:500, AF2864; R&D Systems); guinea pig anti-tdT (RFP) (1:1500,390-004; Synaptic systems); rabbit anti-RFP (1:1500,600-401-379; Rockland); rabbit anti-Timp1 (1:800, 16644-1-AP; Proteintech); sheep anti-Trem2 (1:400, AF1729; Novus); rabbit anti-Tyrobp (1:400,12492S; Cell Signaling); rat anti-Vim (1:200, MAB2105; Novus). All antibodies used were sourced from commercial vendors and were selected because they had previously been validated for fluorescent immunohistochemistry (IHC) in mouse tissue and had manufacturer provided demonstration of specificity based on Western Blots and in most cases validation in peer-reviewed publications.

*Fluorescence secondary antibodies* Alexa 488 (green) or Cy3 (550, red) or Alexa 647, (far red), all from (Jackson Immunoresearch Laboratories, USA). Mouse primary antibodies were visualized using the Mouse-on-Mouse detection kit (M.O.M., Vector). Nuclear stain: 4′,6′-diamidino-2-phenylindole dihydrochloride (DAPI, blue; 2 ng ml−1; Molecular Probes, USA). Sections were cover-slipped using ProLong Gold anti-fade reagent (Invitrogen, Grand Island, NY). Sections were examined and photographed with an epifluorescence microscope using structured illumination hardware and deconvolution software (Zeiss, Oberkochen, Germany).

### Fresh tissue harvest and freezing for RiboTag or snRNAseq

Uninjured mice, or mice at various time points after complete crush SCI, were perfused with ice cold heparinized saline that was prepared using RNAse- and DNAse-free water and 10X PBS for 2 minutes at 7ml/min for blood clearance. Spinal cords were rapidly dissected on ice chilled blocks. For SCI samples, the lesion core center was identified and a tissue block of 1 mm rostral and caudal was then rapidly removed. Anatomically equivalent regions of spinal cord were taken from uninjured mice including from postnatal samples. Tissue samples were rapidly snap frozen in microcentrifuge tubes maintained in a dry ice bath and stored at −80°Cuntil further processing.

### RiboTag immunoprecipitation and RNA-sequencing

Frozen spinal cord tissue was processed by RiboTag Immunoprecipitation (IP) ^31^ using established methods^26,40^. Briefly, tissue was homogenized in RiboTag lysis buffer and centrifuged to remove tissue debris. IP of HA-positive ribosomes was performed by incubating with Anti-HA.11 Epitope Tag Antibody (Biolegend, Cat# 901515) for 4 hours in a microcentrifuge tube on a microtube rotator kept at 4°C. IP solutions were combined with Pierce A/G Magnetic Beads (Thermofisher, #PI88803) and incubated overnight on a microtube rotator at 4°C. On the second day, the solution was separated from the magnetic beads and processed as the “flow through” sample representing mRNA of other cells not from RiboTag-positive cells. Magnetic beads were washed three times with high salt solution (50 mM Tris pH 7.4, 300 mM KCl, 12 mM MgCl2, 1% NP-40, 1 mM Dithiothreitol (DTT), 100 mg/ml Cyclohexamide). Unpurified RNA was collected from the magnetic beads by addition of RLT Plus buffer with BME and vigorous vortexing. RNA was then purified using RNeasy Plus Mini (for *in vitro* cell pellets) or Micro Kits (for spinal cord tissue) (QIAGEN Cat# 74134 and 74034). Total mRNA derived from the RiboTag IP was quantified using a 2100 Bioanalyzer (Agilent) with RNA samples having an RNA integrity numbers (RIN) greater than seven being processed for RNA-Sequencing. Sequencing was performed on poly-A selected libraries using Illumina NovaSeq S2 (UCLA Technology Center for Genomics & Bioinformatics) using pair end reads (2×50 – 50bp length) with an average of 50-100M reads per sample split over two lanes of the S2 flow cell.

### Transcriptomics analysis of RiboTag RNA-Sequencing data

Analysis of RNA-Seq raw data was performed in Galaxy using standardized workflows as done previously^40^. The R1 and R2 FASTQ files from lanes 1 and 2 that were obtained directly from the Illumina NovaSeq S2 run were concatenated and cleaned-up using the Trimmomatic tool. Data were then aligned to the M. musculus (mm10) reference genome using the HISAT2 tool applying default parameters. Gene counts from the aligned datasets was performed using the featureCounts tool applying default parameters. Fragments Per Kilobase of transcript per Million mapped reads (FPKM) values were calculated for each gene directly in Excel (Microsoft) using standardized lists of gene lengths and normalization of the count data. Differential expressed gene (DEG) analysis on raw gene count data was conducted using Edge-R in Galaxy applying Benjamini and Hochberg p-value adjustment and TMM normalization. Across all studies we used a conservative false discovery rate (FDR) cut off < 0.01 to define significance of DEGs and evaluated at least 4 unique samples per experimental group. Gene ontology analyses were performed using Enrichr tool (https://maayanlab.cloud/Enrichr/). Differences in transcript expression across samples were evaluated using data-dimensionality reduction techniques including Principal Component Analysis and Euclidian distance as described below. Heatmaps of DEG data were generated using NG-CHM BUILDER (https://build.ngchm.net/NGCHM-web-builder/). Violin plots of DEGs were generated using Prism 9.

### Threshold criteria for gene expression in healthy astrocytes

Healthy astrocyte expressed genes (AEGs) were defined as having an FPKM value greater than 0.1, which was a conservative cut-off that accounted for genes that were no more than one standard deviation below the mean of the log transformed dataset and represented the upper 77% of all genes that had detectable counts in the dataset. Genes with an FPKM value at or above this threshold in healthy astrocytes were considered healthy expressed genes (“EGs”).

### Isolation and precipitation of astrocyte nuclei for snRNAseq

Using fresh frozen spinal cord tissue harvested as described above, astrocyte-enriched nuclei were isolated by Sox9 antibody-binding and magnet-assisted nuclear immunoprecipitation (MAN-IP) using well characterized procedures^74–76^.To avoid potential batch effects, frozen tissue samples were collected from entire experiments and were then processed at the same time. Tissue samples were first gently dissociated by trituration and pelleted by centrifugation. Nuclei were extracted from cell pellets by gentle resuspension in ice-cold lysis buffer (10mM Tris buffer, 10mM NaCl, 3 mM MgCl2, 0.1% Nonidet P40 Substitute). Nuclei were pelleted by centrifugation (500RPM (Model 5415R, Eppendorf) for 5 min at 4°C) and then resuspended in Nuclei Wash and Resuspension Buffer (NWRB) (1xPBS, 1% BSA, 0.2U/uL RNAse inhibitor) before being washed once more in NWRB and then filtered using a 5 mL Polystyrene Round-Bottom Tube with 35 µm Cell-Strainer Cap and concentrated to a nuclei concentration of 1000 nuclei/µL (1 x 10^6^ nuclei/mL), resuspended and incubated with Sox9 rabbit monoclonal antibody (ThermoFisher, Cat#72016) for 30 minute and then centrifuged at 700g for 10 minutes. The pellet was resuspended in 80μl of MACS Buffer composed of 1X PBS (Tissue Culture grade; Ca^2+^, Mg^2+^ free), 0.5% Nuclease free Bovine Serum Albumin (BSA), and 2mM EDTA, and then incubated with anti-Rabbit IgG Microbeads (Miltenyi, Cat# 130-048-602) for 20 minutes at 4°C. After washing with 1ml of MACS buffer at 300g for 10 minutes at 4 °C, immunolabeled nuclei were enriched by magnetic separation using MACS MS columns (Miltenyi Cat# 30-042-201, 130-042-102 and 130-042-303). The pellet was resuspended in an appropriate volume of Resuspension Buffer to achieve a final concentration of 1000 nuclei/µL (1 x 10^6^ nuclei/mL) and immediately processed using Chromium Next GEM Single Cell 3’ v3.1 kits using manufacturer’s instructions. Libraries processed for RNA-Sequencing using Illumina NovaSeq S4 (housed in the UCLA Technology Center for Genomics & Bioinformatics) using pair end reads (2×100).

### Analysis of single Nuclei RNA-Seq data

Raw single nuclei sequencing data was processed using RNA StarSolo on the Galaxy Single Cell Omics platform using M. musculus (mm10) reference genome, 3M-february-2018 barcode whitelist, the gencode vM25 annotation list, and Cell Ranger v3 configure chemistry options. Scanpy tools were used through the Galaxy platform to convert genes, barcodes and matrix files derived from RNA StarSolo into an AnnData matrix h5ad format. Downstream analysis was performed in R (v4.1) using Seurat package (v4). The dataset was pre-processed to remove cells with high mitochondrial reads, filter out genes that were detected in less than 200 cells and filter out low quality cells that had less than 500 attributed genes. Next, we log2 normalized the dataset, identified the 4000 most variable genes across the total population, computed principal components and summarized the top 50 principal components using the UMAP projection. Cell clustering was performed on the UMAP data via Seurat using Louvain clustering algorithms with a resolution of 0.3 which resulted in 20 discrete clusters for nuclei from both uninjured and SCI tissue. Clusters were annotated as enriched for Astrocytes, Neurons, Oligodendrocytes, Oligodendrocyte Precursor Cells (OPC), or Microglia on the basis of panels of multiple cell-type specific marker genes. Custers identified as astrocyte-enriched were extracted for re-clustering and further extraction based on cohorts of multiple astrocyte specific marker genes, which confidently identified 15,637 astrocyte nuclei that were used for final analyses.

### Principal Component Analysis (PCA)

Principal Component Analysis (PCA) and Euclidean distance analysis was performed using XLStat (Addinsoft Inc, Long Island City, NY) ^40^. For PCA presented throughout, the first two principal components were used to display 2 dimensional scatterplots. PCA factor analysis to identify genes correlated to a particular PC used a factor loading (FL) threshold of > |0.8|. PCA data was represented as Euclidean distance plots throughout. Euclidean distance magnitude calculations were derived by assessing the vector magnitude in PCA space of a specific sample referenced to another sample as an initial point.

### Statistics, power calculations, group sizes and reproducibility

Graph generation and statistical evaluations of repeated measures were conducted by one-way or two-way ANOVA with post hoc independent pair wise analysis as per Tukey, or by Student’s t-tests where appropriate using Prism 10 (GraphPad Software Inc, San Diego, CA). Statistical details of experiments can be found in the figure legends including the statistical tests used and the number of replicative samples. Across all statistical tests significance was defined as p-value <0.05. Power calculations to determine group sizes were performed using G*Power Software V 3.1.9.2. For immunohistochemical quantification analysis and RNA-Sequencing, group sizes were calculated to provide at least 80% power when using the following parameters: probability of type I error (alpha) = .05, a conservative effect size of 0.25, 2-5 treatment groups with multiple measurements obtained per replicate. All graphs show mean values plus or minus standard error of the means (S.E.M.) as well as individual values as dot plots. All bar graphs are overlaid with dot plots where each dot represents the value for one animal to show the distribution of data and the number (N) of animals per group. Injections of NPC and hydrogel formulations were repeated independently at least three times in different colonies of mice across a two-year period with similar results.

## Data availability

Raw FASTQ sequencing files and processed count data have been deposited at Gene Expression Omnibus (GEO) and are publicly available with Accession Number GSE241628. All data generated for this study are included in the main and supplementary figures. For all quantitative figures, files of statistics source data are provided with the paper. Other data that support the findings of this study are available on reasonable request from the corresponding authors.

### Acknowledgements

This work was supported by the Dr. Miriam and Sheldon G. Adelson Medical Foundation (M.V.S., R.K.); US National Institutes of Health (NS084030 to M.V.S.); Paralyzed Veterans of America Research Foundation (T.M.O’S.); Wings for Life (M.V.S. and T.M.O’S.), and Microscopy Core Resource of UCLA Broad Stem Cell Research Center.

## Author contributions

T.M.O’S. and M.V.S. conceptualized and led the overall project. T.M.O’S. and M.V.S. designed, guided and supervised all experiments. T.M.O’S. and S.W. performed surgeries. T.M.O’S., Y.A., and A.C., conducted histological processing. T.M.O’S., and S.W., conducted biochemical and transcriptome processing. T.M.O’S, Y.A., S.W., R.K., V.S., and M.V.S. analyzed data. T.M.O’S. and M.V.S. wrote the manuscript, with input from all co-authors.

## Extended Data Figures

**Extended Data Fig. 1.**
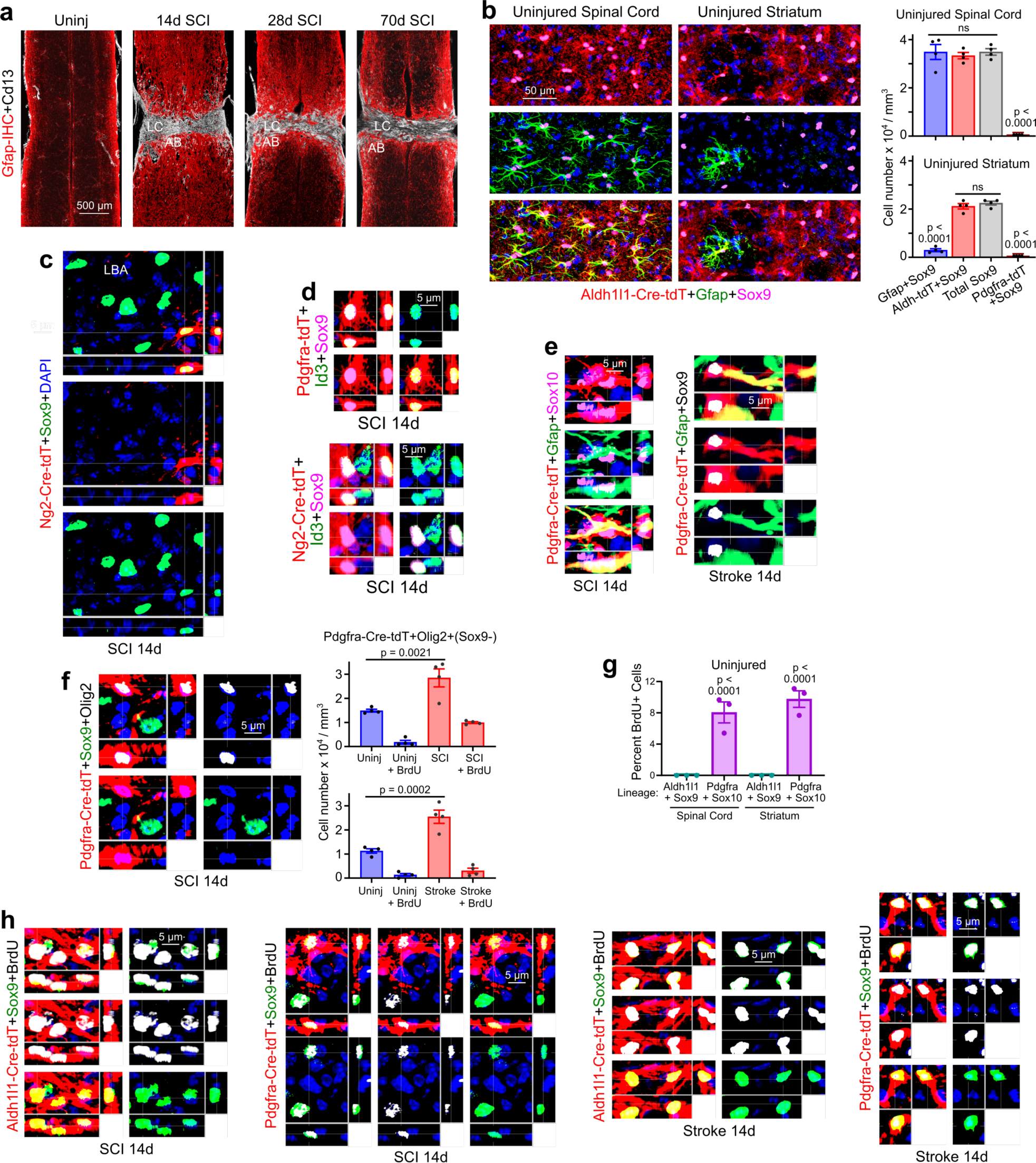
CreERT-tdT lineage tracing of border-forming astrocytes that surround CNS lesions. **a.** Survey images of horizontal sections of spinal cord uninjured and at progressive times after SCI stained by immunohistochemistry (IHC) for astrocytes (Gfap) or stromal cells (CD13), showing maturation and persistence of the astrocyte border (AB) over time. **b.** IHC and cell counts show that in uninjured spinal cord essentially all Aldh1l1-Cre-tdT labelled astrocytes also express Gfap and Sox9, whereas in uninjured striatum only about 13% do. In healthy spinal cord or striatum, no Gfap+ and Sox9+ astrocytes also detectably express Pdgfra-CreERT-tdT. **c.** NG2-Cre-tdT-positive OPC-lineage-derived Sox9-positive lesion border astrocyte (LBA) plus many Sox9-positive but NG2-Cre-tdT-negative Sox9-positive LBA after SCI. **d.** Examples of Pdgfra-Cre-tdT-positive NG2-Cre-tdT-positive OPC-lineage-derived LBA that express the transcription factor Id3. **e.** Examples of Pdgfra-Cre-tdT-positive OPC-lineage-derived LBA that express Gfap, Sox10, and Sox9. **f.** IHC and cell counts show that Pdgfra-Cre-tdT-positive OPCs that express Olig2 but not Sox9 are also present in the lesion border zone and that a portion of these OPCs proliferate and are labeled with BrdU after SCI or stroke. **g.** Cell counts show that in healthy cord or striatum there are no astrocytes detectably labelled with BrdU and dividing during a 6-day delivery period whereas, about 8 to 10% of OPC were BrdU-labelled and newly proliferated during the same delivery period. **h.** Examples of Aldh1l1-Cre-tdT-positive or Pdgfra-Cre-tdT-positive newly proliferated LBA that express Sox9 and are labelled with BrdU after SCI or stroke.

**Extended Data Fig. 2.**
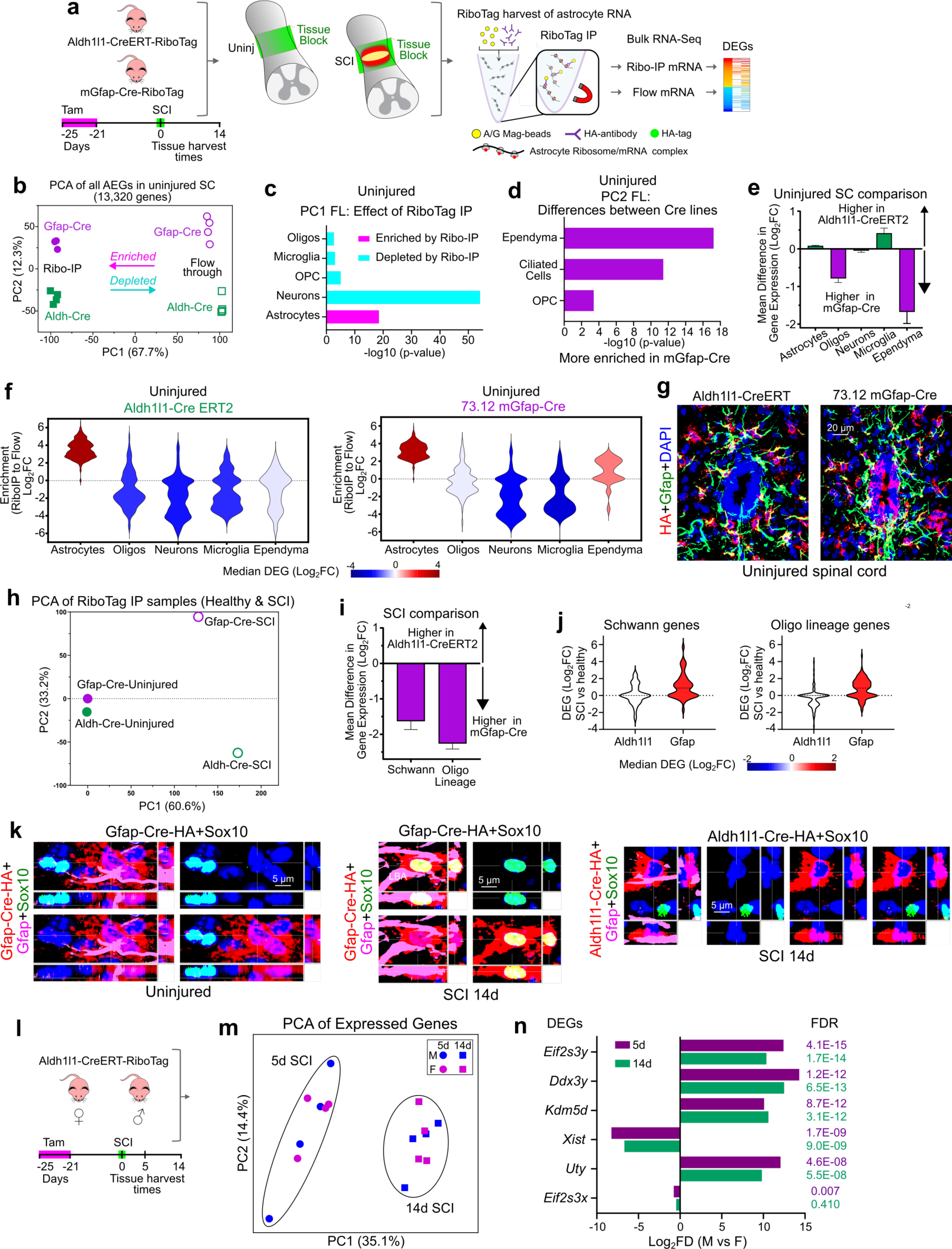
Comparison of transcriptional profiles obtained with tamoxifen-activated Aldh1l1-Cre-ERT-RiboTag or constitutive mGfap-Cre-RiboTag, and comparison of astrocyte transcriptional responses to SCI in female or male mice. **a.** Experimental design for bulk RNAseq of astrocyte expressed genes (EGs) obtained from uninjured mice by using transgenically targeted RiboTag using two different Cre lines, Ald1h1-CreERT or mGfap-Cre. **b.** Principal component analysis (PCA) of all EGs obtained by RNAseq of RiboTag immunoprecipitation (IP) of uninjured astrocyte EGs or flow through of non-astrocyte EGs from all other cells using the two different Cre lines. Note that both Cre lines show similar enrichment levels of astrocyte EGs by RiboTag-IP as demonstrated by PC1 which shows differences in astrocyte EGs relative to flow through EGs. Small differences between the two Cre lines are reflected in PC2. **c.** Cell type analysis of EGs whose factor loading (FC) determined PC1 demonstrates significant and substantial de-enrichment of genes associated with neurons, oligodendrocyte lineage cells and microglia using both Cre lines. Thus, in healthy spinal cord tissue, RNAseq of Aldh1l1-CreERT-RiboTag and mGfap-Cre-RiboTag gave similar transcriptional profiles that were highly significantly enriched for known astrocyte RNA transcripts and de-enriched for known transcripts associated with neurons, oligodendrocyte lineages or microglia. **d.** Cell type analysis of EGs whose FC determined PC2 indicates that differences between the two Cre lines was due to less de-enrichment of genes associated with ependyma, ciliated cells and OPC when using mGfap-Cre-RiboTag. **e,f.** Direct analysis of gene panels known to be associated with specific cell types showed pronounced and equal enrichment of astrocyte EGs and pronounced and essentially equal de-enrichment of neuronal and microglial EGs in both Cre lines, with greater de-enrichment of oligodendrocyte lineage genes using Aldh1l1-CreERT-RiboTag. Notably, ependymal genes were somewhat enriched in mGfap-Cre-RiboTag IPs. These differences could be explained by the expression of Gfap by multipotent radial glial progenitors that give rise to ependyma during development^77^. **g.** As expected, immunohistochemistry demonstrated Gfap-Cre-RiboTag (hemagglutinin, HA) staining in many ependyma cells, which derive from Gfap-positive radial glia during development and express Cre for this reason. Notably, HA was not immunohistochemically detectable in ependyma in our Aldh1l1-CreERT-RiboTag mice. **h.** PCA of RiboTag-IP RNAseq of uninjured astrocyte EGs or astrocytes after SCI comparing the two different Cre lines. Note that SCI induced similar levels of changes in astrocyte transcriptional profiles measured using the two Cre lines, as as demonstrated by PC1 of astrocyte DEGs. Differences between the two Cre lines are reflected in PC2. **i,j.** Cell type analysis of DEGs whose factor loading (FC) determined PC2 demonstrates significantly higher expression after SCI of DEGs associated with Schwann cells and oligodendrocyte lineage cells when using the mGfap-Cre line. This could be explained by the constitutive expression of mGfap-Cre-RiboTag and the acquisition after SCI of Gfap expression either by astrocytes derived from OPCs or by OPC-derived Schwann cells^32,78^. **k.** IHC for RiboTag (HA) shows that in uninjured mice, astrocytes that expressed mGfap-Cre-HA and Gfap protein are not co-labeled with Sox10, whereas after SCI, some mGfap-Cre-HA and Gfap-protein expressing cells astrocytes also express Sox10. **l.** Experimental design comparing SCI-induced transcriptional changes in female and male mice using the Aldh1l1-CreERT-RiboTag approach. **m.** PCA with unbiased clustering shows that transcriptional profiles cluster according to time after SCI with no detectable sex differences. **n.** Only six Y-chromosome genes exhibited sex differences in expression by astrocytes after SCI.

**Extended Data Fig. 3.**
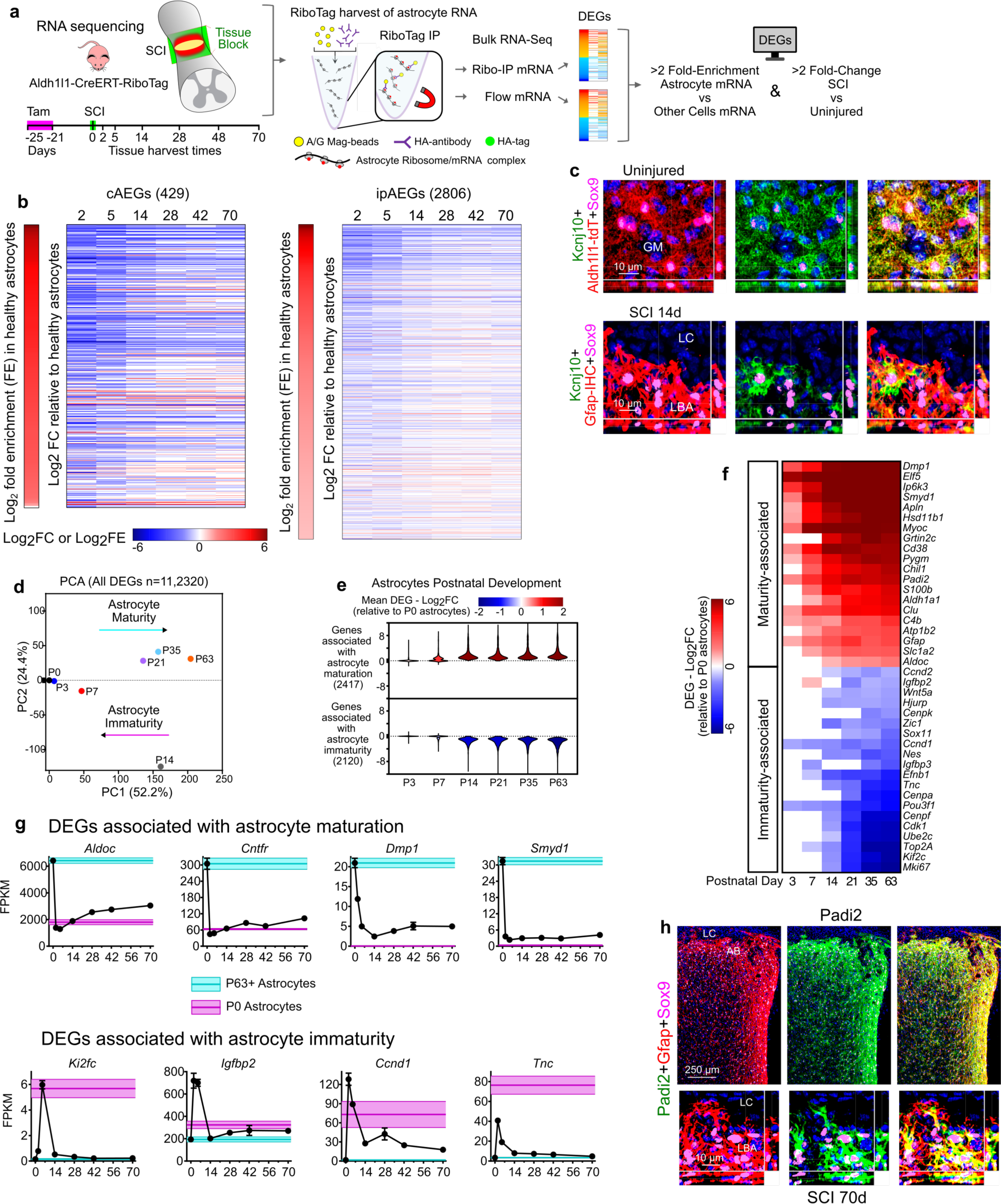
SCI-induced astrocyte dedifferentiation and proliferation, and expression profiles of genes associated with astrocyte immaturity or astrocyte maturity during development and after SCI. **a.** Experimental design for enrichment analysis of gene expression by astrocytes compared with other cells by comparing FPKM levels of astrocyte RNA precipitated via RiboTag with FPKM levels of RNA from other cells collected in non-precipitated flow through eluates. **b.** Heatmaps of log2 fold changes (Log_2_FC) in gene expression over time after SCI of 429 cAEGs and 2806 ipAEGs enriched in astrocytes by at least 2-fold and shown as log2 fold enrichment (Log_2_FE) compared with other cells. We confirmed that expression of all 429 cAEGs identified from the literature were enriched in our samples from health astrocytes by at least 2-fold and up to over 50-fold compared with non-astrocyte genes. **c.** Immunohistochemistry shows markedly reduced Kcnj10 (Kir4.1) protein immunoreactivity in lesion border astrocytes (LBA) compared with astrocytes in uninjured grey matter (GM). **d.** PCA of all DEGs showing DEG cohorts detected at various postnatal days (P) and thereby defining specific timepoints. **e.** Mean changes in genes associated with astrocyte maturity or immaturity at different postnatal (P) days. **f.** Heatmap of Log_2_FC changes in gene expression over time of selected genes associated with astrocyte maturity or immaturity at different postnatal (P0 or P63) days. **g.** Time course of expression changes (FPKM, fragments per kilobase of transcript per million reads) after SCI of selected genes associated with astrocyte maturity or immaturity compared with mean expression levels at different P0 or P63 days as markers of immature or mature levels, respectively. **h.** Immunohistochemistry of Padi2 protein at 70 days after SCI, an astrocyte maturity gene that is down regulated acutely after SCI and then returns to normal expression levels (see main Fig. 3m) and becomes prominently expressed in lesion border astrocytes (LBA).

**Extended Data Fig. 4.**
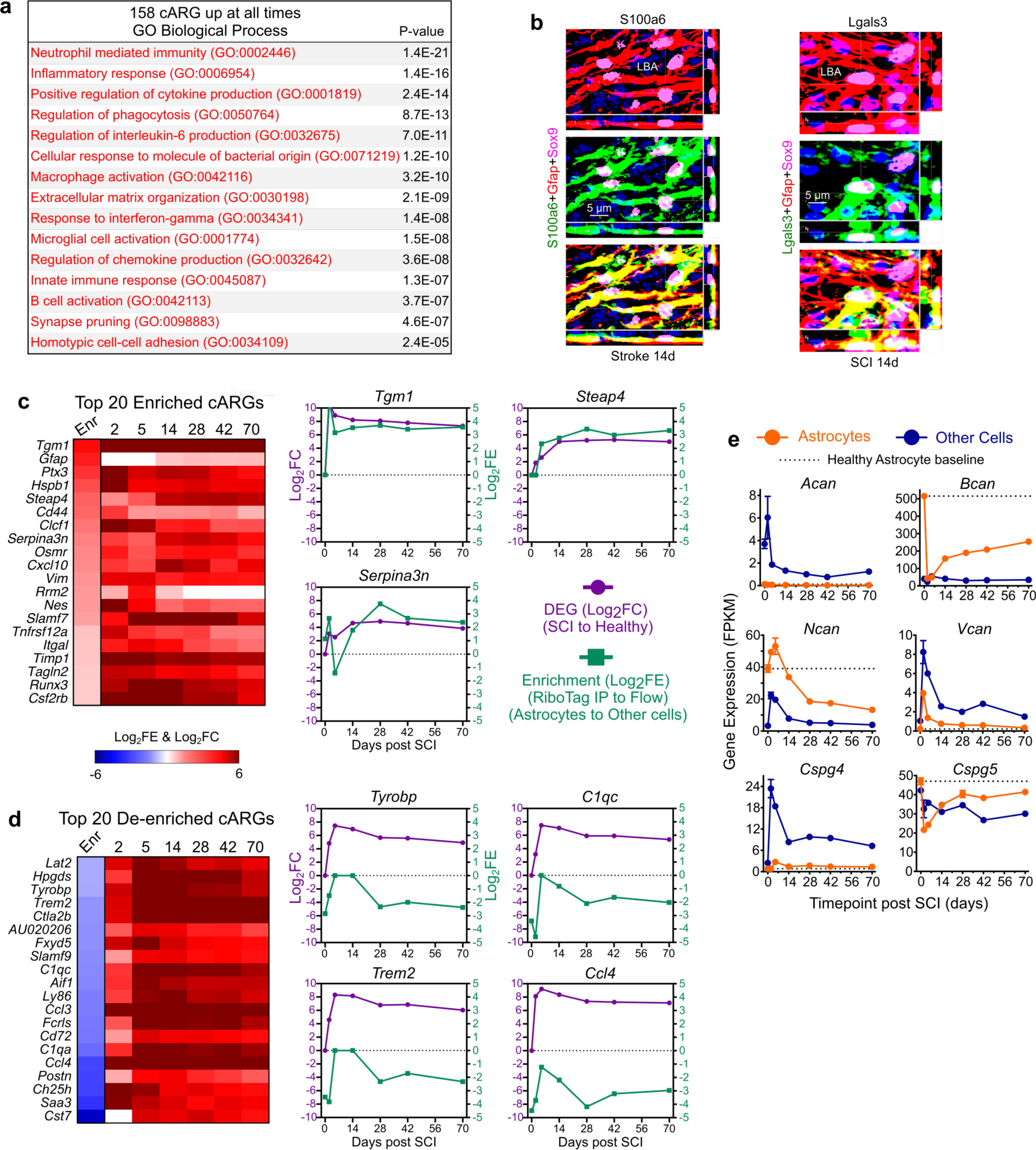
Reactivity DEGs. **a.** Top GO Biological Processes associated with 158 cARGs upregulated at all times after SCI. **b.** Immunohistochemistry of S100a6 and Lgals3 protein in lesion border astrocytes (LBA) after stroke (S100a6) or SCI (Lgals3). Note that while S100a6 is present in the processes of most LBA, Lgals3 is present in some but not others. **c,d.** Heatmaps of mean log_2_FC after SCI of 20 cARGs that are either most enriched (**c**) or most de-enriched (**d**) in astrocytes compared with other cells; graphs compare log_2_FC with mean log_2_FE in selected examples. **e.** Expression (FPKM) by astrocytes and other cells of chondroitin sulphate proteoglycans (CSPGs) after SCI. Note that the prototypical CSPG, *Acan*, is not detectably expressed by astrocytes at any time after SCI, and *Vcan* and *Csp4* were far more highly expressed by non-astrocytes. Only *Bcan* and *Ncan* are more highly expressed by astrocytes, but *Bcan* expression declines after SCI to levels below healthy, and although *Ncan* expression increases at 5 days after SCI, by 28 days it also declined to levels lower than in healthy astrocytes.

**Extended Data Fig. 5.**
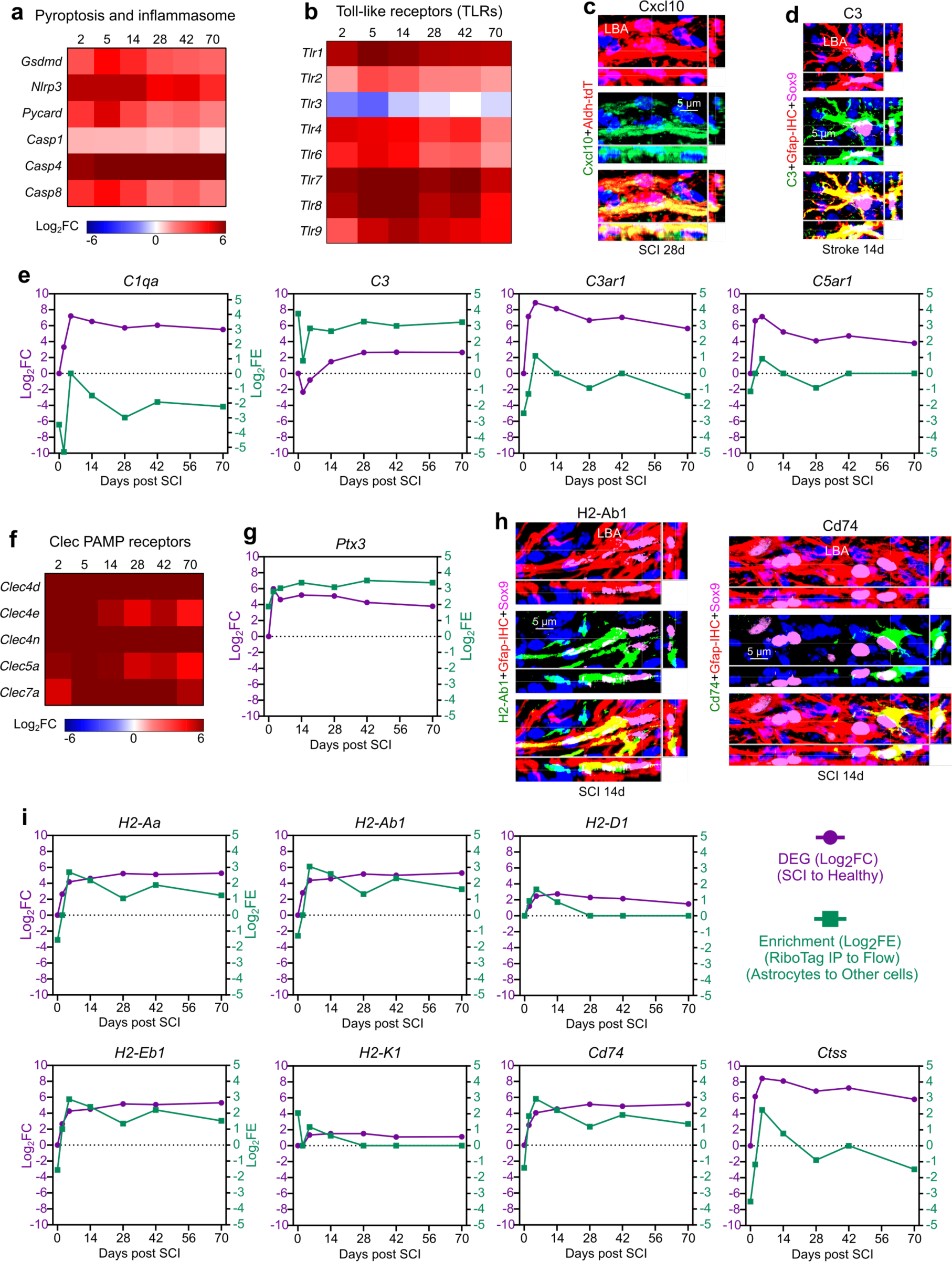
Innate and adaptive immune regulatory signaling, and contributions to antimicrobial defense and antigen presentation. **a,b.** Heatmaps of Log_2_FC of DEGs associated with astrocyte pyroptosis and inflammasome generation (**a**) or Toll-like receptors (**b**) at different days after SCI. **c,d.** Immunohistochemistry of Cxcl10 protein after SCI (**c**) and C3 protein after stroke in lesion border astrocytes (LBA). (**d**). **e.** Graphs compare changes in log_2_FC with changes in mean log_2_FE after SCI of selected complement related DEGs. **f.** Heatmaps of Log_2_FC of *Clec* DEGs encoding pathogen-associated molecular patterns (PAMPs) receptors. **g.** Graph showing both upregulation of expression (log_2_FC) and enrichment (log_2_FE) in border astrocytes of antimicrobial *Pitx3*. **h.** Immunohistochemistry showing intense expression of H2-Ab1 protein or C74 protein in scattered border astrocytes while other nearby astrocytes do not exhibit detectable levels. **i.** Graphs compare changes in log_2_FC with changes in mean log_2_FE after SCI of selected DEGs associated with antigen presentation.

**Extended Data Fig. 6.**
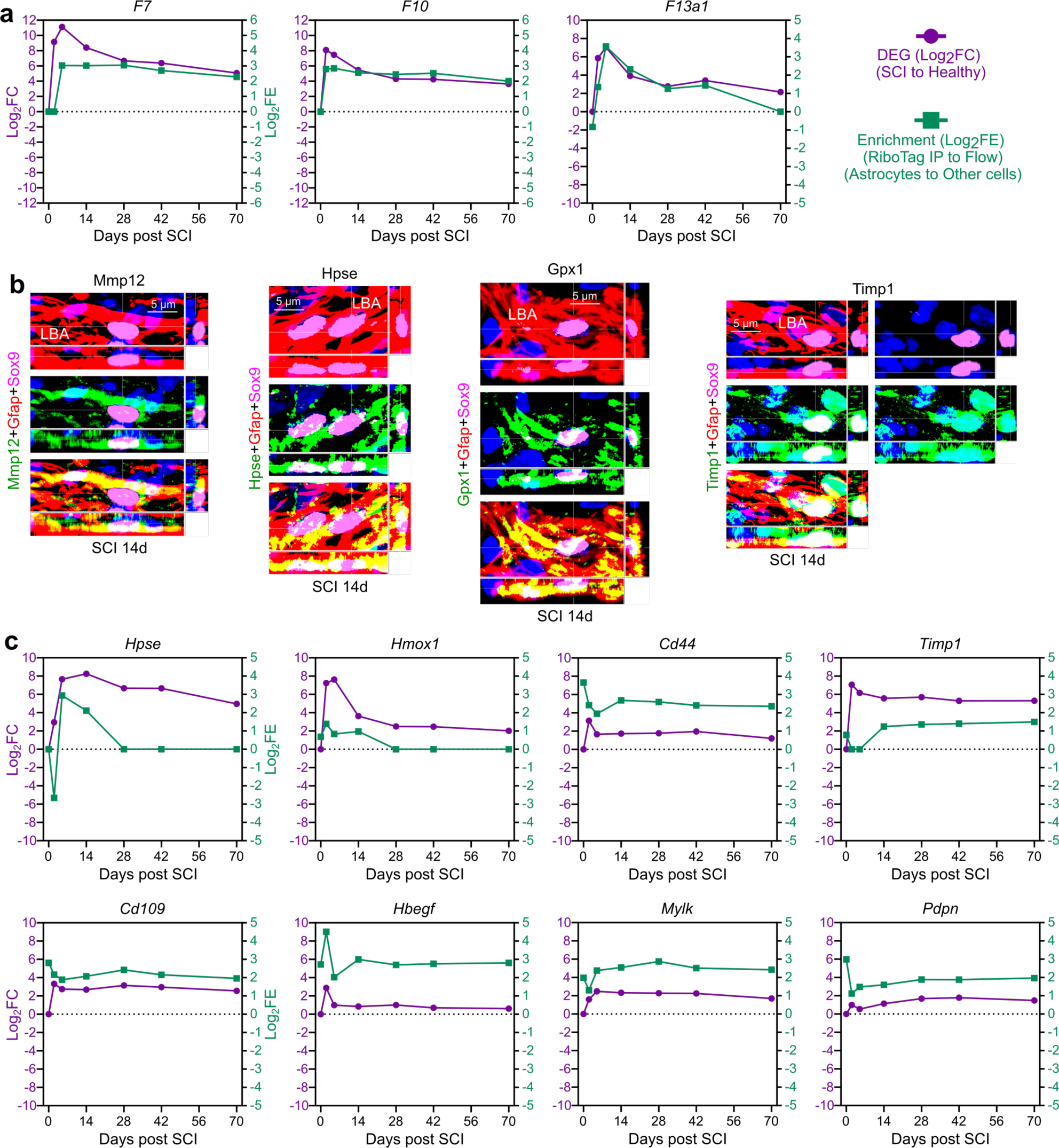
Wound healing associated DEGs. **a.** Graphs compare changes in log_2_FC with changes in mean log_2_FE after SCI of selected coagulation factor associated DEGs. **b.** Immunohistochemistry of Mmp12, Hpse, Gpx1 and Timp1 proteins in lesion border astrocytes (LBA) after SCI. **c.** Graphs compare changes in log_2_FC with changes in mean log_2_FE after SCI of selected diverse wound healing associated DEGs.

**Extended Data Fig. 7.**
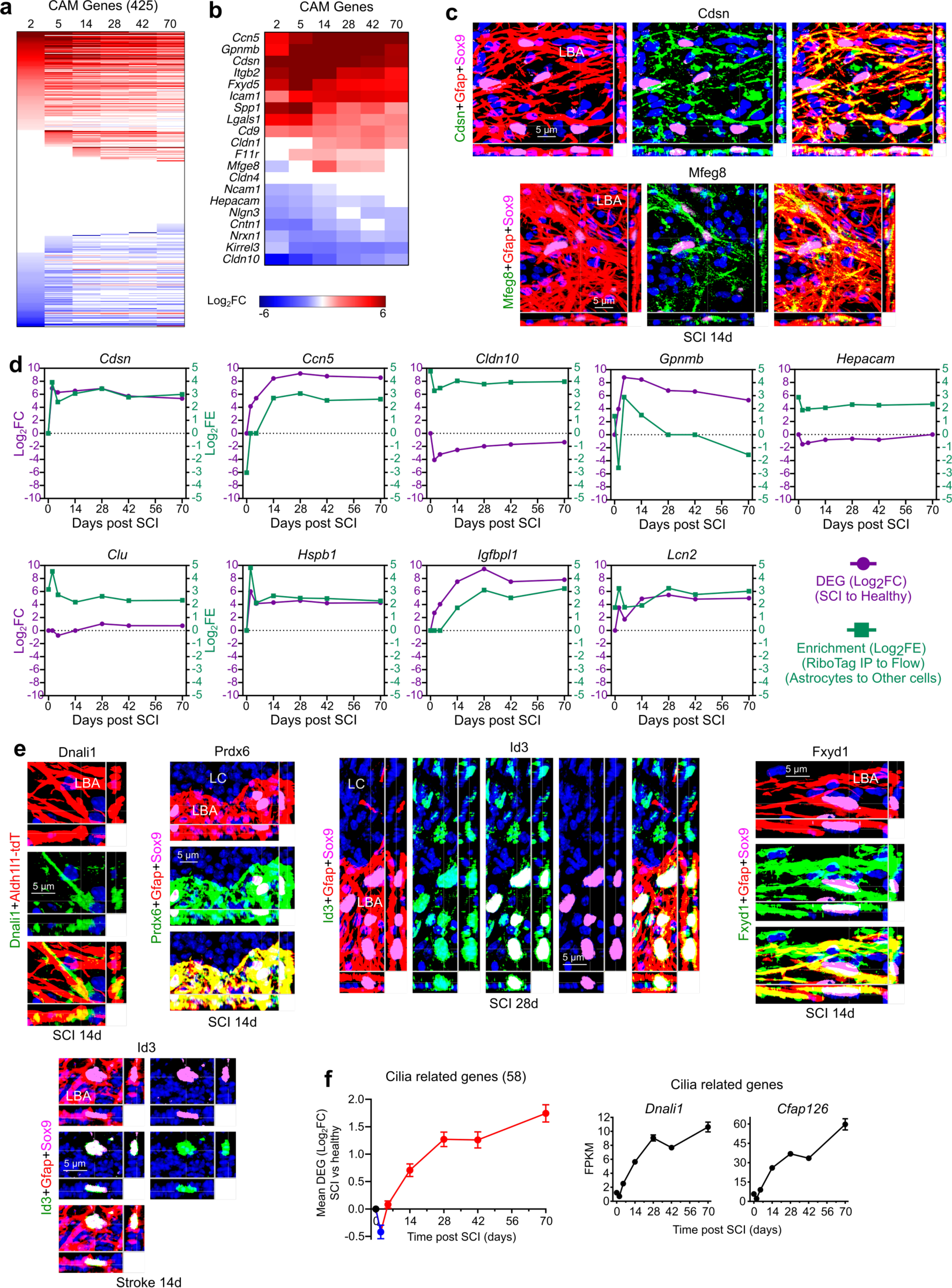
Cell adhesion molecules (CAMs) and delayed astrocyte reactivity DEGs (dARGs). **a,b.** Heatmaps of Log_2_FC of all (**a**) and selected (**b**) CAM-associated DEGs at different times after SCI. **c.** Immunohistochemistry of Cdsn and Mfeg6 CAM proteins in lesion border astrocytes (LBA) after SCI. **d.** Graphs comparing changes in log_2_FC with changes in mean log_2_FE after SCI of selected dARGs. **e.** Immunohistochemistry of Dnali1, Prdx6, Id3 and Fxyd1 proteins in LBA after SCI or stroke. **f.** Graphs show delayed mean upregulation after SCI of 58 cilia related DEGs and representative examples of *Dnali1* and *Cfap126*.

**Extended Data Fig. 8.**
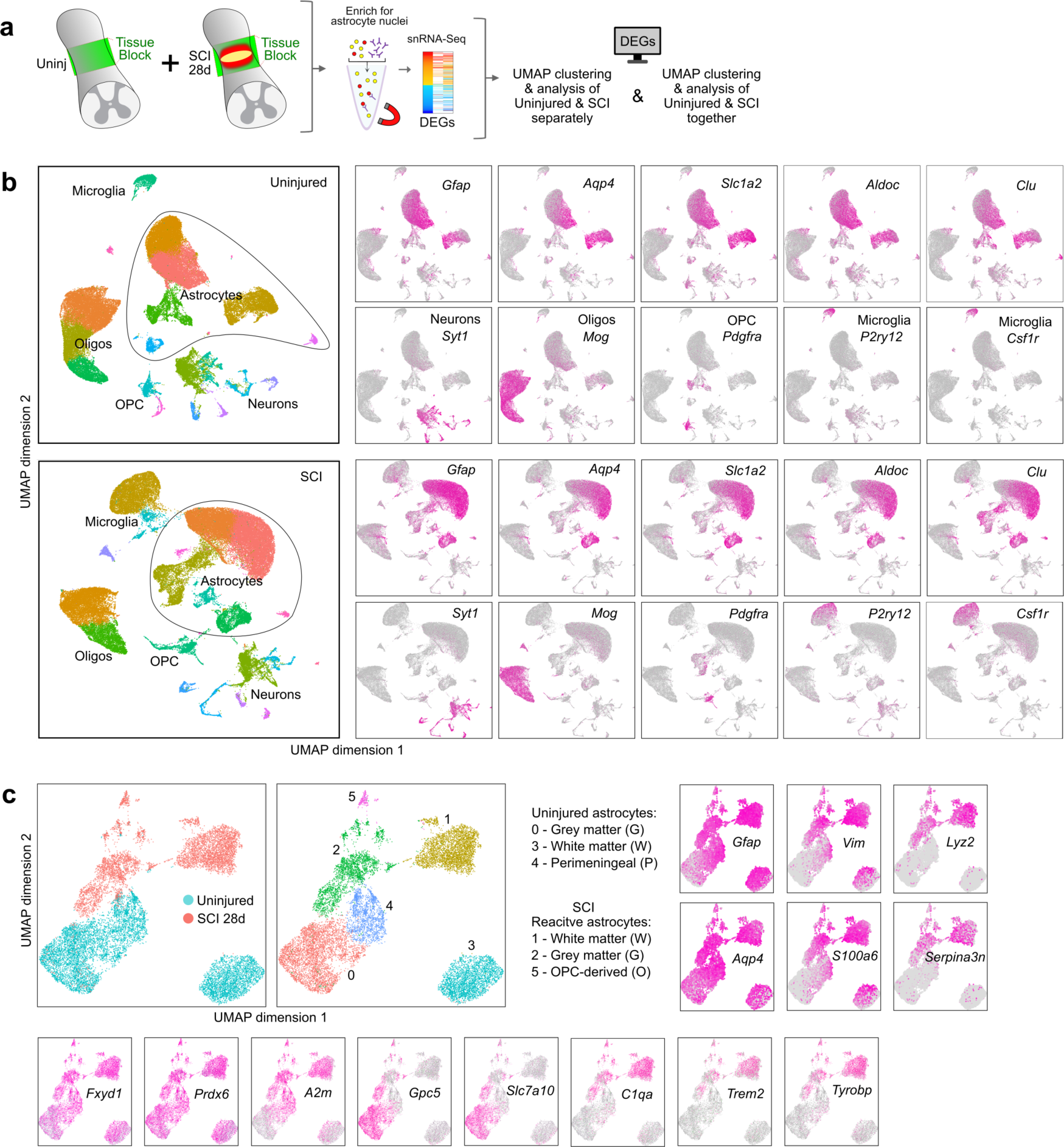
Single nucleus RNA sequencing (snRNAseq). **a.** Experimental design for snRNAseq of the same region of thoracic spinal cord tissue from uninjured mice and mice at 28 days after SCI. **b.** UMAP clusters of all nuclei. Clusters were annotated as enriched for Astrocytes, Neurons, Oligodendrocytes (Oligos), Oligodendrocyte Precursor Cells (OPC), or Microglia based on panels of multiple cell-type specific marker genes. Custers identified as astrocyte-enriched were extracted for re-clustering and further extraction based on cohorts of multiple astrocyte specific marker genes, which confidently identified 15,637 astrocyte nuclei that were used for final analyses as shown in main figure 7. **c.** UMAP clusters of astrocyte nuclei and feature plots of selected DEGs.

**Extended Data Fig. 9.**
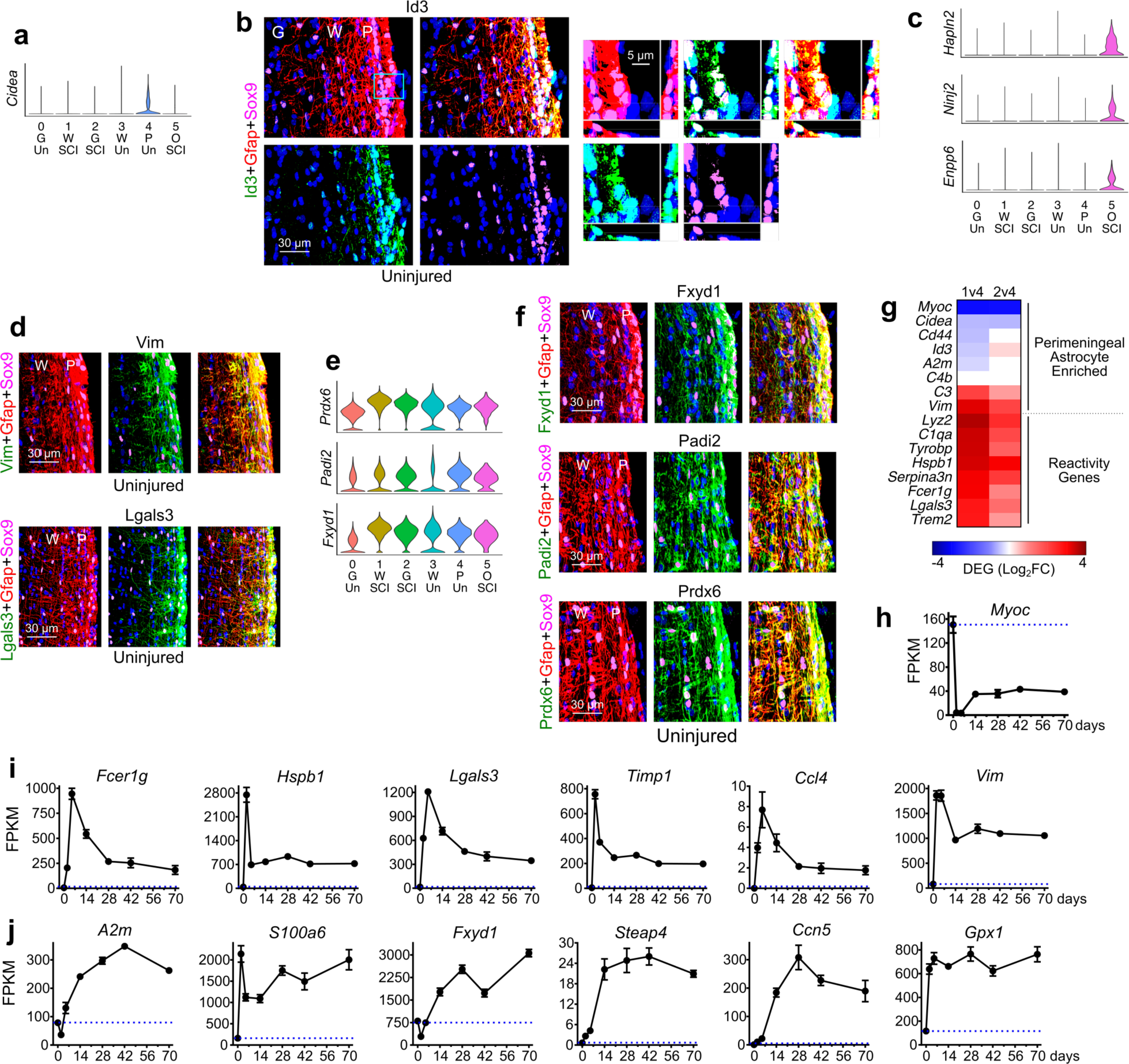
Molecular characteristics of perimeningeal astrocytes (PMA) and mature lesion border astrocytes (LBA). **a.** Violin plot of *Cidea* expression limited to cluster 4 of uninjured PMA (P). **b.** Immunohistochemistry shows Id3 protein levels are high in PMA (P), low in white matter astrocytes (W) and low or not detectable in grey matter astrocytes (G). **c.** Violin plots show OPC lineage markers *Hapln2, Ninj2, Enpp6* expression limited to cluster 5 of OPC-derived LBA (O). **d.** Immunohistochemistry shows Vim and Lgals3 protein levels are highly enriched in PMA (P). **e.** Violin plots show enriched expression of shows *Fxyd1*, *Padi2* and *Prdx6* in reactive astrocyte clusters (1,2,5) and PMA (5) compared with uninjured grey (0) or white (3) matter astrocytes. **f.** Immunohistochemistry shows Fxyd1, Padi2 and Prdx6 protein levels are enriched in PMA (P) relative to white matter astrocytes. **g.** Heatmap comparing expression levels in cluster 1 (reactive astrocytes) with cluster 4 (uninjured PMA) of DEGs enriched in PMA (top) or astrocyte reactivity DEGs (bottom). **h.** Astro-RiboTag RNAseq shows an acute and persisting decline in *Myoc* expression after SCI. **i,j.** Additional selected astrocyte DEGs exhibiting different patterns of expression changes in the form of acute rise followed by decline (i) or delayed but persistent increase (j) after SCI as detected by Astro-RiboTag RNAseq.

**Extended Data Fig. 10.**
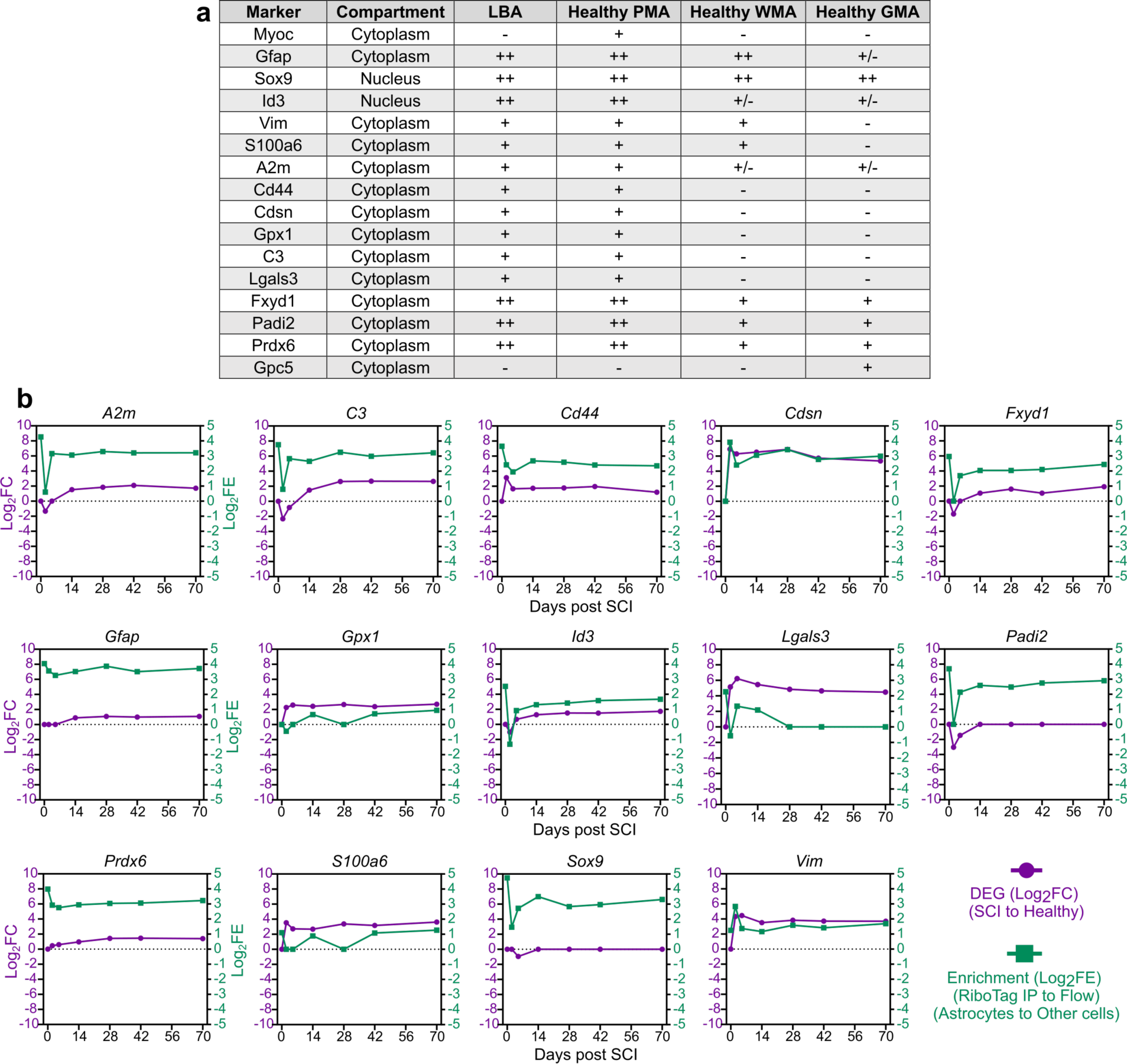
Comparisons of protein immunohistochemistry, DEG changes over time after SCI and DEG enrichment in astrocytes compared with other cells, for various astrocyte associated molecules, including potential markers enriched in mature border-forming wound repair astrocytes. **a.** Semi-quantitative (++, +, +/−, or −) observer scored comparison of immunoreactivity levels of various molecular markers across lesion border astrocytes (LBA) and astrocytes in healthy CNS tissue, perimeningeal astrocytes (PMA), grey matter astrocytes (GMA) and white matter astrocytes (WMA). **b.** Graphs compare changes in log_2_FC with changes in mean log_2_FE after SCI of DEGs encoding the molecular markers of LBA shown in a. Note that even though DEG expression of potential markers may be enriched in LBA compared with other cells, protein expression is in many cases not exclusive to LBA, and protein is often expressed also by adjacent non-neural cells in lesion cores, such as for C3, Lgals3, S100a6, Vim and others. Thus, these markers are not exclusive to LBA, and for this reason, co-detection of these markers together with Gfap and Sox9 as done here is essential to reliably identify LBA.

## References (Main text)

1. Gurtner, G.C., Werner, S., Barrandon, Y. & Longaker, M.T. Wound repair and regeneration. Nature 453, 314–321 (2008).

2. Sofroniew, M.V. & Vinters, H.V. Astrocytes: biology and pathology. Acta Neuropathol 119, 7–35 (2010).

3. Burda, J.E. & Sofroniew, M.V. Reactive gliosis and the multicellular response to CNS damage and disease. Neuron 81, 229–248 (2014).

4. Sofroniew, M.V. Astrocyte barriers to neurotoxic inflammation. Nat Rev Neurosci 16, 249–263 (2015).

5. Linnerbauer, M., Wheeler, M.A. & Quintana, F.J. Astrocyte Crosstalk in CNS Inflammation. Neuron 108, 608–622 (2020).

6. Escartin, C., et al. Reactive astrocyte nomenclature, definitions, and future directions. Nat Neurosci 24, 312–325 (2021).

7. Gage, F.H. Mammalian neural stem cells. Science 287, 1433–1438 (2000).

8. Jessen, N.A., Munk, A.S., Lundgaard, I. & Nedergaard, M. The Glymphatic System: A Beginner’s Guide. Neurochem Res 40, 2583–2599 (2015).

9. Khakh, B.S. & Sofroniew, M.V. Diversity of astrocyte functions and phenotypes in neural circuits. Nat Neurosci 18, 942–952 (2015).

10. Allen, N.J. & Eroglu, C. Cell Biology of Astrocyte-Synapse Interactions. Neuron 96, 697–708 (2017).

11. Verkhratsky, A. & Nedergaard, M. Physiology of Astroglia. Physiol Rev 98, 239–389 (2018).

12. Khakh, B.S. & Deneen, B. The Emerging Nature of Astrocyte Diversity. Annu Rev Neurosci 42, 187–207 (2019).

13. Wanner, I.B., et al. Glial scar borders are formed by newly proliferated, elongated astrocytes that interact to corral inflammatory and fibrotic cells via STAT3-dependent mechanisms after spinal cord injury. J Neurosci 33, 12870–12886 (2013).

14. Marescal, O. & Cheeseman, I.M. Cellular Mechanisms and Regulation of Quiescence. Developmental cell 55, 259–271 (2020).

15. Sofroniew, M.V. Astrocyte Reactivity: Subtypes, States, and Functions in CNS Innate Immunity. Trends Immunol 41, 758–770 (2020).

16. Burda, J.E., et al. Divergent transcriptional regulation of astrocyte reactivity across disorders. Nature 606, 557–564 (2022).

17. Bush, T.G., et al. Leukocyte infiltration, neuronal degeneration, and neurite outgrowth after ablation of scar-forming, reactive astrocytes in adult transgenic mice. Neuron 23, 297–308 (1999).

18. Buffo, A., et al. Origin and progeny of reactive gliosis: A source of multipotent cells in the injured brain. Proc. Natl. Acad. Sci. USA 105, 3581–3586 (2008).

19. Bardehle, S., et al. Live imaging of astrocyte responses to acute injury reveals selective juxtavascular proliferation. Nat Neurosci 16, 580–586 (2013).

20. Sirko, S., et al. Astrocyte reactivity after brain injury-: The role of galectins 1 and 3. Glia 63, 2340–2361 (2015).

21. O’Shea, T.M., Burda, J.E. & Sofroniew, M.V. Cell biology of spinal cord injury and repair. J Clin Invest 127, 3259–3270 (2017).

22. Chen, M., et al. Leucine Zipper-Bearing Kinase Is a Critical Regulator of Astrocyte Reactivity in the Adult Mammalian CNS. Cell reports 22, 3587–3597 (2018).

23. Frik, J., et al. Cross-talk between monocyte invasion and astrocyte proliferation regulates scarring in brain injury. Embo Rep 19(2018).

24. O’Shea, T.M., et al. Foreign body responses in mouse central nervous system mimic natural wound responses and alter biomaterial functions. Nature communications 11, 6203 (2020).

25. Faulkner, J.R., et al. Reactive astrocytes protect tissue and preserve function after spinal cord injury. J. Neurosci. 24, 2143–2155 (2004).

26. Anderson, M.A., et al. Astrocyte scar formation aids central nervous system axon regeneration. Nature 532, 195–200 (2016).

27. Williamson, M.R., Fuertes, C.J.A., Dunn, A.K., Drew, M.R. & Jones, T.A. Reactive astrocytes facilitate vascular repair and remodeling after stroke. Cell reports 35, 109048 (2021).

28. Lahiri, A., et al. Astrocytic deletion of protein kinase R-like ER kinase (PERK) does not affect learning and memory in aged mice but worsens outcome from experimental stroke. J Neurosci Res (2023).

29. Kretzschmar, K. & Watt, F.M. Lineage tracing. Cell 148, 33–45 (2012).

30. Ren, Y., et al. Ependymal cell contribution to scar formation after spinal cord injury is minimal, local and dependent on direct ependymal injury. Sci Rep 7, 41122 (2017).

31. Sanz, E., et al. Cell-type-specific isolation of ribosome-associated mRNA from complex tissues. Proc Natl Acad Sci U S A 106, 13939–13944 (2009).

32. Zawadzka, M., et al. CNS-resident glial progenitor/stem cells produce Schwann cells as well as oligodendrocytes during repair of CNS demyelination. Cell Stem Cell 6, 578–590 (2010).

33. Sozmen, E.G., et al. Nogo receptor blockade overcomes remyelination failure after white matter stroke and stimulates functional recovery in aged mice. Proc Natl Acad Sci U S A 113, E8453–E8462 (2016).

34. Hackett, A.R., et al. Injury type-dependent differentiation of NG2 glia into heterogeneous astrocytes. Experimental Neurology 308, 72–79 (2018).

35. Hesp, Z.C., et al. Proliferating NG2-Cell-Dependent Angiogenesis and Scar Formation Alter Axon Growth and Functional Recovery After Spinal Cord Injury in Mice. J Neurosci 38, 1366–1382 (2018).

36. Srinivasan, R., et al. New Transgenic Mouse Lines for Selectively Targeting Astrocytes and Studying Calcium Signals in Astrocyte Processes In Situ and In Vivo. Neuron 92, 1181–1195 (2016).

37. Kang, S.H., Fukaya, M., Yang, J.K., Rothstein, J.D. & Bergles, D.E. NG2+ CNS glial progenitors remain committed to the oligodendrocyte lineage in postnatal life and following neurodegeneration. Neuron 68, 668–681 (2010).

38. Zhu, X., Bergles, D.E. & Nishiyama, A. NG2 cells generate both oligodendrocytes and gray matter astrocytes. Development 135, 145–157 (2008).

39. Zhu, X., et al. Age-dependent fate and lineage restriction of single NG2 cells. Development 138, 745–753 (2011).

40. O’Shea, T.M., et al. Lesion environments direct transplanted neural progenitors towards a wound repair astroglial phenotype in mice. Nature communications 13, 5702 (2022).

41. Hernandez, V.G., et al. Translatome analysis reveals microglia and astrocytes to be distinct regulators of inflammation in the hyperacute and acute phases after stroke. Glia (2023).

42. Baldwin, K.T., et al. HepaCAM controls astrocyte self-organization and coupling. Neuron (2021).

43. Allen, N.J., et al. Astrocyte glypicans 4 and 6 promote formation of excitatory synapses via GluA1 AMPA receptors. Nature 486, 410–414 (2012).

44. Jopling, C., Boue, S. & Izpisua Belmonte, J.C. Dedifferentiation, transdifferentiation and reprogramming: three routes to regeneration. Nature reviews. Molecular cell biology 12, 79–89 (2011).

45. Gotz, M., Sirko, S., Beckers, J. & Irmler, M. Reactive astrocytes as neural stem or progenitor cells: In vivo lineage, In vitro potential, and Genome-wide expression analysis. Glia 63, 1452–1468 (2015).

46. Moonen, S., et al. Pyroptosis in Alzheimer’s disease: cell type-specific activation in microglia, astrocytes and neurons. Acta Neuropathol (2022).

47. Sun, H., et al. Bacteria reduce flagellin synthesis to evade microglia-astrocyte-driven immunity in the brain. Cell reports 40, 111033 (2022).

48. Vivinetto, A.L., et al. Zeb2 Is a Regulator of Astrogliosis and Functional Recovery after CNS Injury. Cell reports 31, 107834 (2020).

49. Klatt Shaw, D., et al. Localized EMT reprograms glial progenitors to promote spinal cord repair. Developmental cell 56, 613–626 e617 (2021).

50. Marconi, G.D., et al. Epithelial-Mesenchymal Transition (EMT): The Type-2 EMT in Wound Healing, Tissue Regeneration and Organ Fibrosis. Cells 10(2021).

51. Dongre, A. & Weinberg, R.A. New insights into the mechanisms of epithelial-mesenchymal transition and implications for cancer. Nature reviews. Molecular cell biology 20, 69–84 (2019).

52. Bohrer, C., et al. The balance of Id3 and E47 determines neural stem/precursor cell differentiation into astrocytes. The EMBO journal 34, 2804–2819 (2015).

53. Venugopal, N., et al. The primary cilium dampens proliferative signaling and represses a G2/M transcriptional network in quiescent myoblasts. BMC Mol Cell Biol 21, 25 (2020).

54. Urban, N., Blomfield, I.M. & Guillemot, F. Quiescence of Adult Mammalian Neural Stem Cells: A Highly Regulated Rest. Neuron 104, 834–848 (2019).

55. Wu, Y.E., Pan, L., Zuo, Y., Li, X. & Hong, W. Detecting Activated Cell Populations Using Single-Cell RNA-Seq. Neuron 96, 313–329 e316 (2017).

56. Wei, H., et al. Glial progenitor heterogeneity and key regulators revealed by single-cell RNA sequencing provide insight to regeneration in spinal cord injury. Cell reports 42, 112486 (2023).

57. Berry, M., et al. Deposition of scar tissue in the central nervous system. Acta Neurochir Suppl (Wien) 32, 31–53 (1983).

58. Bunge, R.P., Puckett, W.R. & Hiester, E.D. Observations on the pathology of several types of human spinal cord injury, with emphasis on the astrocyte response to penetrating injuries. Advances in neurology 72, 305–315 (1997).

59. Norenberg, M.D., Smith, J. & Marcillo, A. The pathology of human spinal cord injury: defining the problems. J Neurotrauma 21, 429–440 (2004).

60. Paolicelli, R.C., et al. Microglia states and nomenclature: A field at its crossroads. Neuron 110, 3458–3483 (2022).

61. Konishi, H. & Kiyama, H. Microglial TREM2/DAP12 Signaling: A Double-Edged Sword in Neural Diseases. Frontiers in cellular neuroscience 12, 206 (2018).

62. Clarke, E.V. & Tenner, A.J. Complement modulation of T cell immune responses during homeostasis and disease. J Leukoc Biol 96, 745–756 (2014).

63. Dai, D.L., Li, M. & Lee, E.B. Human Alzheimer’s disease reactive astrocytes exhibit a loss of homeostastic gene expression. Acta neuropathologica communications 11, 127 (2023).

64. Askenazi, M., et al. Compilation of reported protein changes in the brain in Alzheimer’s disease. Nature communications 14, 4466 (2023).

65. Clark, I.C., et al. Barcoded viral tracing of single-cell interactions in central nervous system inflammation. Science 372(2021).

66. Barnabe-Heider, F., et al. Origin of new glial cells in intact and injured adult spinal cord. Cell Stem Cell 7, 470–482 (2010).

67. Muthusamy, N., Brumm, A., Zhang, X., Carmichael, S.T. & Ghashghaei, H.T. Foxj1 expressing ependymal cells do not contribute new cells to sites of injury or stroke in the mouse forebrain. Sci Rep 8, 1766 (2018).

68. Rhett, J.M., et al. Novel therapies for scar reduction and regenerative healing of skin wounds. Trends Biotechnol 26, 173–180 (2008).

69. Rog-Zielinska, E.A., Norris, R.A., Kohl, P. & Markwald, R. The Living Scar--Cardiac Fibroblasts and the Injured Heart. Trends in molecular medicine 22, 99–114 (2016).

70. Iismaa, S.E., et al. Comparative regenerative mechanisms across different mammalian tissues. NPJ Regen Med 3, 6 (2018).

71. Li, Y., et al. Microglia-organized scar-free spinal cord repair in neonatal mice. Nature 587, 613–618 (2020).

72. Maxwell, W.L., Follows, R., Ashhurst, D.E. & Berry, M. The response of the cerebral hemisphere of the rat to injury. II. The neonatal rat. Philosophical transactions of the Royal Society of London. Series B, Biological sciences 328, 501–513 (1990).

## References (Methods)

73. Garcia, A.D.R., Doan, N.B., Imura, T., Bush, T.G. & Sofroniew, M.V. GFAP-expressing progenitors are the principle source of constitutive neurogenesis in adult mouse forebrain. Nature Neurosci. 7, 1233–1241 (2004).

74. Krishnaswami, S.R., et al. Using single nuclei for RNA-seq to capture the transcriptome of postmortem neurons. Nature protocols 11, 499–524 (2016).

75. Bhattacharyya, S., Sathe, A.A., Bhakta, M., Xing, C. & Munshi, N.V. PAN-INTACT enables direct isolation of lineage-specific nuclei from fibrous tissues. PLoS One 14, e0214677 (2019).

76. Batiuk, M.Y., et al. An immunoaffinity-based method for isolating ultrapure adult astrocytes based on ATP1B2 targeting by the ACSA-2 antibody. J Biol Chem 292, 8874–8891 (2017).

77. Redmond, S.A., et al. Development of Ependymal and Postnatal Neural Stem Cells and Their Origin from a Common Embryonic Progenitor. Cell reports 27, 429–441 e423 (2019).

78. Assinck, P., et al. Myelinogenic Plasticity of Oligodendrocyte Precursor Cells following Spinal Cord Contusion Injury. J Neurosci 37, 8635–8654 (2017).

